# Human iPSC-derived CNS and retinal microvasculature-on-a-chip models recapitulate hallmarks of diabetic microvascular pathology

**DOI:** 10.64898/2026.01.15.699708

**Authors:** Mareike Volz, Alexander Kleger, Stefan Liebau, Natalia Pashkovskaia

## Abstract

The central nervous system microvasculature (CNS-mv) protects neural tissue from harmful substances while supplying oxygen, nutrients, and signaling molecules. Metabolic diseases such as diabetes induce pathological changes in these microvasculatures, leading to severe outcomes including stroke and diabetic retinopathy. Blood vessels in the brain and retina share similarities and undergo comparable pathological alterations in diabetic patients. However, mechanistic understanding of CNS-mv pathogenesis remains limited due to the lack of physiologically relevant *in vitro* models. Here, we developed three complementary human iPSC-derived microvascular models: (1) a CNS-mv-on-a-chip for mechanistic studies, (2) scalable 3D microvasculature drops for high-throughput screening, and (3) an inner blood-retinal barrier-on-a-chip incorporating Müller glia. All platforms self-assembled into perfusable networks with high pericyte (PC) coverage and barrier function. We identified the TNF-α/NF-κB pathway as the central mediator of hyperglycemia-induced vascular damage, accompanied by robust induction of inflammatory cytokines IL-1β and IL-6. Importantly, cell-type-specific analysis revealed distinct inflammatory roles: endothelial cells (ECs) predominantly activated TNF-α/NF-κB downstream signaling, whereas pericytes selectively upregulated IL-1β, suggesting a coordinated EC-PC inflammatory crosstalk driving vascular pathology. Functionally, both hyperglycemia and TNF-α/IL-1β exposure induced vascular regression, reduced PC coverage, and increased ghost vessel formation, confirming these cytokines as key effectors of diabetic microvascular damage. Uniquely, Müller glia showed divergent behavior from PCs, enhancing their perivascular sheath under inflammation – indicating reactive gliosis rather than protection. Together, these *in vitro* platforms provide a versatile and physiologically relevant hiPSC-based system for mechanistic studies, screening anti-inflammatory therapeutics and developing glia-targeted interventions.

## Introduction

Worldwide, the number of diabetic patients and, accordingly, the prevalence of related vascular complications, including higher risks for ischemic stroke, small vessel disease (SVD), microangiopathy, and diabetic retinopathy (DR), is increasing. Despite decades of research, therapeutic approaches remain largely limited to symptom relief and late-stage disease interventions. Understanding the mechanisms linking metabolic dysfunction to vascular diseases is essential for improving diagnostics and developing effective therapies.

The microvasculature is one of the main parts of the cardiovascular system that plays an essential role in the oxygen and nutrient supply as well as in the delivery of signaling molecules, such as hormones. Endothelial cells (ECs) are the main building blocks of blood vessels and form the wall that regulates the permeability of the vessel. ECs are supported by mural cells: smooth muscle cells serve as the predominant supporting cell type in arterioles and venules, while pericytes (PCs) predominate in the capillaries [1, 2]. PCs play a particularly crucial role in capillary formation in the central nervous system (CNS), where PC density is highest compared to other tissues [3, 4]. The high PCs coverage is essential for maintaining microvascular stability and barrier function, as well as regulation of the cerebral blood flow and immune response [5].

In addition to classical mural cells, CNS microvasculature is supported by glial cells. Astrocytes (ACs) ensheath brain capillaries, while Müller glia cells (MGs) support vessels in the retina [6]. Despite assumed functional similarities, there are no *in vitro* studies comparing both cell types regarding their influence on microvasculature, due to the challenge of obtaining MG from human donors. This knowledge gap is particularly relevant, because capillaries in the retina and brain share similarities. Multiple studies have shown that retinal microvasculature abnormalities and SVD are linked and can be caused by the same processes [7–10]. The resemblance between the cerebral and retinal microvasculature likely reflects their shared embryological origin: their PC populations arise from the neural crest and are therefore neuroectodermal in origin. This shared lineage may result in similar gene expression patterns, making cerebral and retinal PCs morphologically and physiologically related. The association between brain and retinal microvascular disorders supports the use of retinal imaging as a non-invasive method to evaluate ischemic stroke risk in individuals with diabetic conditions [11].

The hyperglycemic and inflammatory milieu disturbs the barrier function of CNS microvasculature [12]. On a molecular level, hyperglycemia triggers oxidative stress and inflammatory response, leading to early pathological changes, including the loss of EC tight junctions, perivascular PC loss, and basement membrane thickening [13–16]. These changes compromise the barrier function and vascular stability. Consequently, this makes the microvasculature fragile and increases vascular permeability, causing fluid accumulation and small hemorrhages. The capillary network becomes narrower due to basement membrane thickening, which in turn reduces the blood flow through vessels [17]. As a compensatory mechanism, to restore blood flow and improve oxygen and nutrient supply, the neovascularization through a VEGF-mediated response is activated [18, 19]. The newly grown blood vessels, however, often show inadequate morphology and pathologically higher permeability, leading to further disease progression.

Current therapies, such as anti-VEGF/ANG2 treatments (bevacizumab, ranibizumab, aflibercept, and faricimab), focus on symptom management rather than addressing the underlying causes of microvascular pathology. Although these therapies have been shown to reduce retinal inflammation [20, 21], they do not target upstream pathogenic mechanisms. Critically, no existing treatments address early-stage disease, when intervention might prevent irreversible neural tissue damage.

The lack of early-disease therapy partly arises from the *in vitro* disease modelling limitations. To date, most studies of microvasculature pathologies caused by diabetes use either animal models or adherent cell culture systems, both of which have significant limitations: rodents do not completely recapitulate the human-specific pathological responses; adherent cell cultures lack complexity, such as the interaction of several cell types and three-dimensional tissue architecture, for accurate disease modeling.

Microfluidic organ-on-chip technology offers an *in vitro* platform for human disease modeling by recreating three-dimensional CNS microvasculature’s architecture and incorporate several cell types, ensuring physiological cellular interactions and functional readouts. Recently microfluidic models recapitulated CNS microvasculature pathology using primary cells [22]. For example, a microvasculature-on-a-chip system reproduced key features of diabetic retinopathy (DR), such as pericyte loss and microvascular degeneration, using primary retinal ECs, PCs, and ACs [23]. However, these models remain limited by challenges associated with sourcing primary human cells. These limitations can be addressed by using human induced pluripotent stem cell (hiPSC)-derived cells. While some studies successfully incorporated hiPSC-derived ECs and ACs [24, 25], none have achieved complete vascular units with hiPSC-derived PCs.

Here, we present the first human iPSC-based microvasculature platform incorporating neural crest-derived PCs. We developed and compared three complementary *in vitro* models: (1) a CNS-mv-on-a-chip for mechanistic studies, (2) scalable microvascular drops for screening, (3) and, to our knowledge, the first inner blood–retinal barrier (iBRB)-on-a-chip incorporating MGs for retina-specific pathology. All three models recapitulated key hallmarks of DR, including vascular regression, reduced PC coverage, and ghost vessel formation. Transcriptomic profiling and vascular morphology analysis revealed TNF-α and IL-1β inflammatory signaling as a primary driver of vascular pathology under hyperglycemic conditions. Cell-type specific gene expression analysis showed that TNF-α signaling was induced in ECs, while PCs amplified the inflammation *via* upregulation of IL1B. The novel iBRB-on-a-chip platform further demonstrated increased MG vessel coverage under inflammatory conditions, suggesting the potential for MG-targeted therapies.

## Results

### hiPSC-derived Endothelial cells and Pericytes form a microvascular network on a chip

To model the central nervous system microvasculature (CNS-mv), we used human induced pluripotent stem cell (hiPSC)-derived endothelial cells (ECs) and neural crest-derived pericytes (PCs) in a microfluidic chip. ECs and PCs were generated following previously published protocols [26, 27] and characterized by the expression of cell-type-specific markers: CD31, VE-cadherin, claudin-5, and ZO-1 for ECs (Supplementary Fig. 1A), and PDGFRβ, NG2, SMA, and desmin for PCs (Supplementary Fig. 1B).

For microvasculature formation, ECs and PCs were loaded into a microfluidic chip at a 1:1 ratio, approximating the physiological condition in the retina and brain [4]. In the microfluidic chip, the cells self-organized into capillary-like structures (Fig. 1A). Within the microvasculature, PCs localized around ECs, providing structural support (Fig. 1B, C, D). PC coverage of the vasculature of 76.1 ± 6.5 % was comparable to physiological levels reported *in vivo* [28, 29]. The formed capillaries possessed both a lumen (Fig. 1B-D) and a basement membrane, as indicated by collagen IV deposition (Fig. 1C-E).

**Figure 1:**
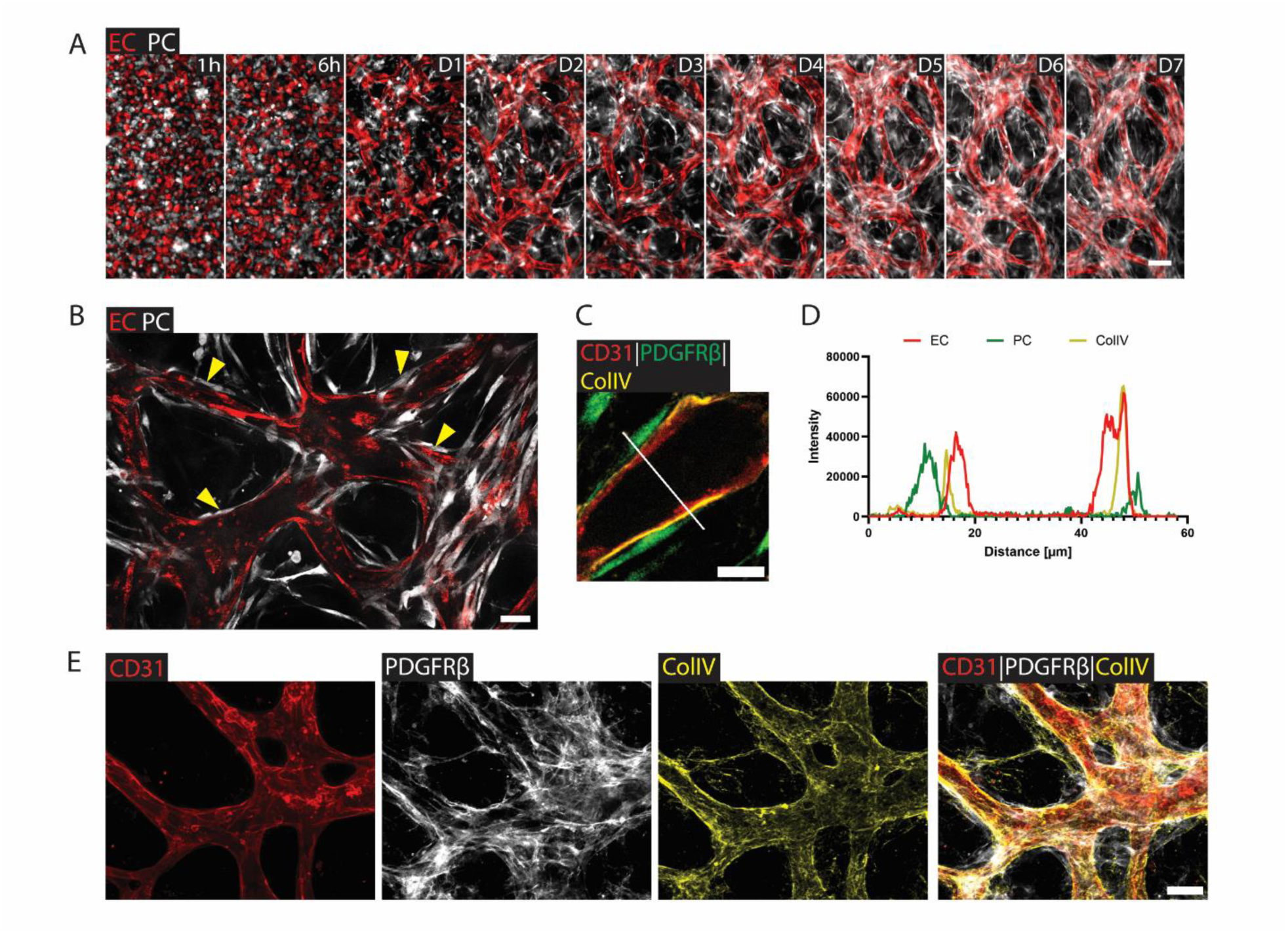
**CNS-mv-on-a-chip formation with hiPSC-derived ECs and PCs.** (A) Fluorescence images show the time course of vessel formation. Endothelial cells (ECs) were labeled with mCherry (red), and pericytes (PCs) with GFP (white). Images were taken at 1 hour, 6 hours, and daily from day 1 to day 7. Scale bar: 100 µm (B) Immunofluorescence image of CNS-mv-on-a-chip with mCherry-labeled ECs (red) and GFP-labeled PCs (white). Yellow arrowheads show PCs covering ECs. Scale bar: 50 µm. (C) Cross-section image of a vessel stained for CD31 (ECs, red), PDGFRβ (PCs, green), and collagen IV (basement membrane, yellow). Scale bar: 20 µm. (D) Plot profile showing the spatial localization of EC, PC, and collagen IV across the vessel section. (E) Immunofluorescence images show CNS-mv-on-a-chip stained for CD31 (ECs, red), PDGFRβ (PCs, white), and collagen IV (basement membrane, yellow). Scale bar: 100 µm.

Together, these results demonstrate that hiPSC-derived ECs and PCs self-organize into a stable, tubular microvascular network within the microfluidic chip.

### Assessment of barrier function in CNS-mv-on-a-chip

The main function of the central nervous system microvasculature (CNS-mv) is to regulate selective transport of oxygen, nutrients, and signaling molecules from the blood to neural tissue while restricting the entry of potentially harmful substances. Key elements responsible for this barrier function are EC junctions, including tight junction proteins (ZO-1 and claudin-5) and adherens junction proteins (VE-cadherin). We confirmed that ECs in the CNS-mv-on-a-chip expressed VE-cadherin, claudin-5, and ZO-1, indicating the presence of both adherens and tight junctions (Fig. 2A).

**Figure 2:**
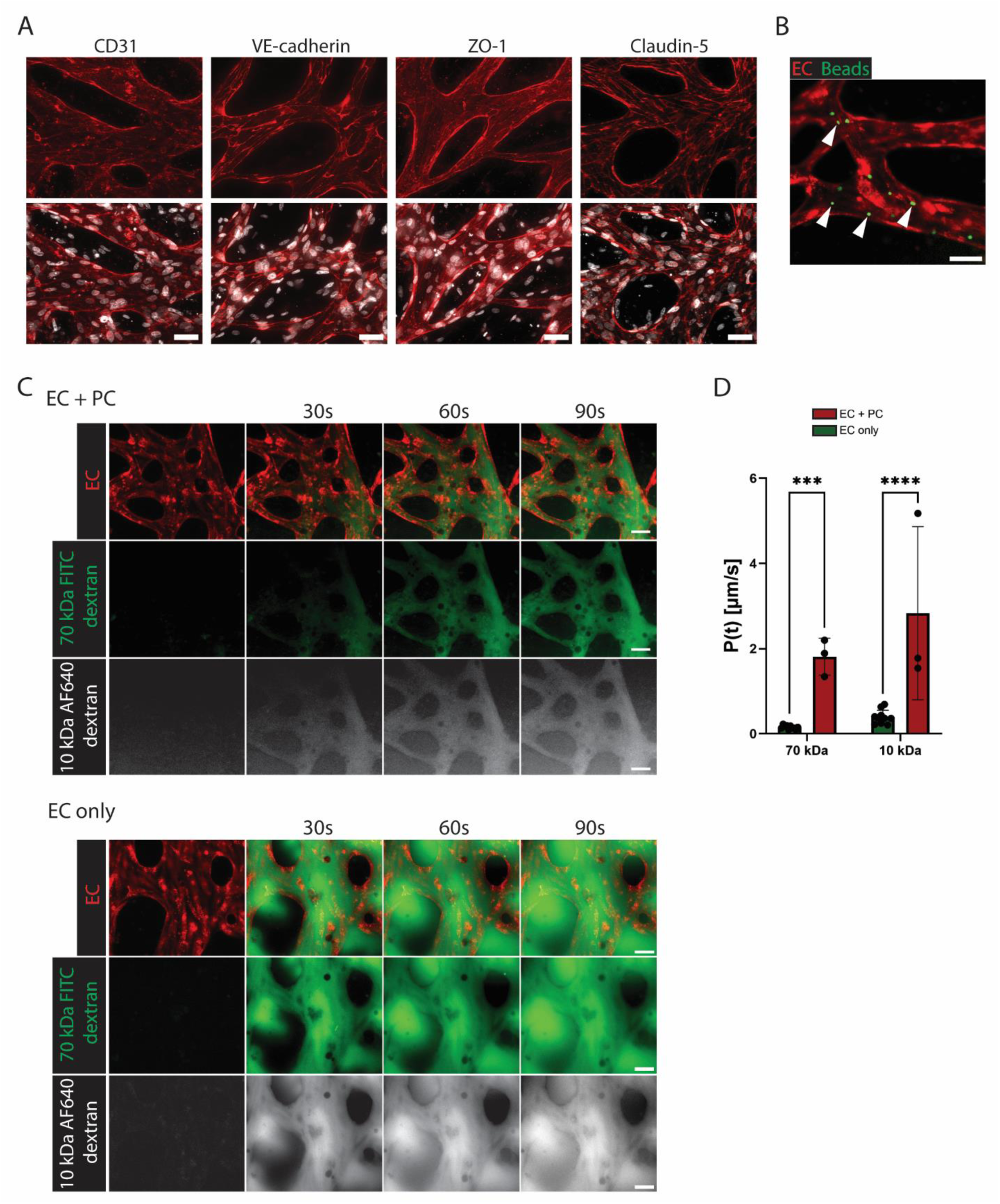
**Analysis of CNS-mv-on-a-chip barrier function.** (A) Immunofluorescence images of CNS-mv-on-a-chip stained for CD31 (ECs), ZO-1 and claudin-5 (tight-junction proteins), and VE-cadherin (adherent junction protein). Scale bar: 50 µm. (B) Fluorescence image of perfused CNS-mv-on-a-chip vessels with AF-488-labeled glass beads (7.5 µm in diameter). ECs were mCherry-labeled. Arrowheads point to perfused beads inside the vessels. Scale bar: 50 µm. (C) Fluorescence images of the CNS-mv-on-a-chip perfusion with 70 kDa FITC-dextran (green) and 10 kDa AF-640-dextran (grey). The top panel shows vasculature formed by ECs and PCs in a 1:1 ratio; the bottom panel EC-alone vasculature. ECs were mCherry-labeled. Scale bar: 50 µm. (D) Quantification of the apparent permeability coefficient of capillaries formed with and without PCs. N = 2-3, n = 3-10.

To evaluate whether the microvascular network contained a functional lumen accessible to circulating blood cells, we perfused the chip with AF-488-labeled glass beads (7.5 µm in diameter), which are comparable in size to lymphocytes (Fig. 2B). The beads flowed through the capillaries and remained confined within the vessel walls formed by mCherry-labeled ECs, demonstrating physiological intraluminal flow and perfusability (Fig. 2B).

We next assessed the barrier function in CNS-mv-on-a-chip using fluorescently labeled dextran molecules of different molecule sizes: 70 kDa FITC-dextran and 10 kDa AF-640-dextran. Under physiological conditions, CNS-mv restricts the passage of macromolecules bigger than 4 kDa [30]. Consistent with this, both dextran molecules remained mostly confined within the lumen (Fig. 2C), with low apparent permeability coefficients [31]: 0.15 ± 0.04 µm/s for 70 kDa and 0.39 ± 0.16 µm/s for 10 kDa (Fig. 2D). Low apparent permeability coefficient values with limited size-dependent molecular leakage, confirm the formation of a restrictive barrier in the CNS-mv-on-a-chip.

Interestingly, although ECs can form vessel-like structures without PC support, they cannot establish a functional barrier on their own. In the absence of PCs, barrier integrity was significantly compromised, with increased permeability coefficients: 1.81 ± 0.43 µm/s for 70 kDa dextran and 2.83 ± 2.03 µm/s for 10 kDa dextran (Fig. 2C, D). These data highlight the essential role of PCs in maintaining CNS-mv barrier function.

In conclusion, our CNS-mv-on-a-chip model recapitulates key features of the CNS-mv barrier, including tight junction formation, functional perfusion, and low macromolecular permeability, validating this platform as a reliable system for studying CNS vascular networks.

### RNA sequencing reveals increased inflammatory cytokine signalling under hyperglycemic conditions

After establishing the CNS-mv-on-a-chip, we next aimed to model diabetic conditions using the platform. As hyperglycemia is a key initiating factor in diabetic pathologies, we exposed CNS-mv-on-a-chip to hyperglycemia for 7 days and analyzed transcriptomic profiles using RNA sequencing.

Principal component analysis (PCA) revealed clear separation between the high-glucose (75 mM glucose) and control (5.5 mM glucose) groups (Fig. 3A, Supplementary Fig. 2B), indicating distinct gene expression profiles. Differential gene expression (DGE) analysis identified 144 upregulated and 119 downregulated genes in the hyperglycemic group. Among the upregulated genes were inflammatory cytokines such as IL1B (2.2 Log2FC) and IL6 (2.6 Log2FC), the matrix-remodeling protease MMP9 (1.2 Log2FC), and PLVAP (1.4 Log2FC), which is associated with increased vascular permeability (Fig, 3B) [32, 33]. Genes linked to altered glucose metabolic stress, including PDK4 (2.1 Log2FC) [34, 35] and ARRDC4 (2.5 Log2FC) [34, 35] were also significantly upregulated, indicating metabolic stress in hyperglycemic CNS-mv-on-a-chip. In contrast, downregulated genes included TRIB3 (−2.4 Log2FC), a negative regulator of NF-κB and modulator of glucose transport, and NTN4 (−1.4 Log2FC), whose reduced expression is associated with vascular destabilization (Fig. 3B, Supplementary Fig. 2C) [36, 37]. Further analysis revealed the robust upregulation of inflammation related genes, including cytokines such as CXCL8 (2.2 Log2FC), CCL2 (1.8 Log2FC), CXCL2 (2.0 Log2FC), CXCL1 (1.9 Log2FC), and IL6 (2.6 Log2FC), consistent with inflammatory response (Fig. 3C).

**Figure 3:**
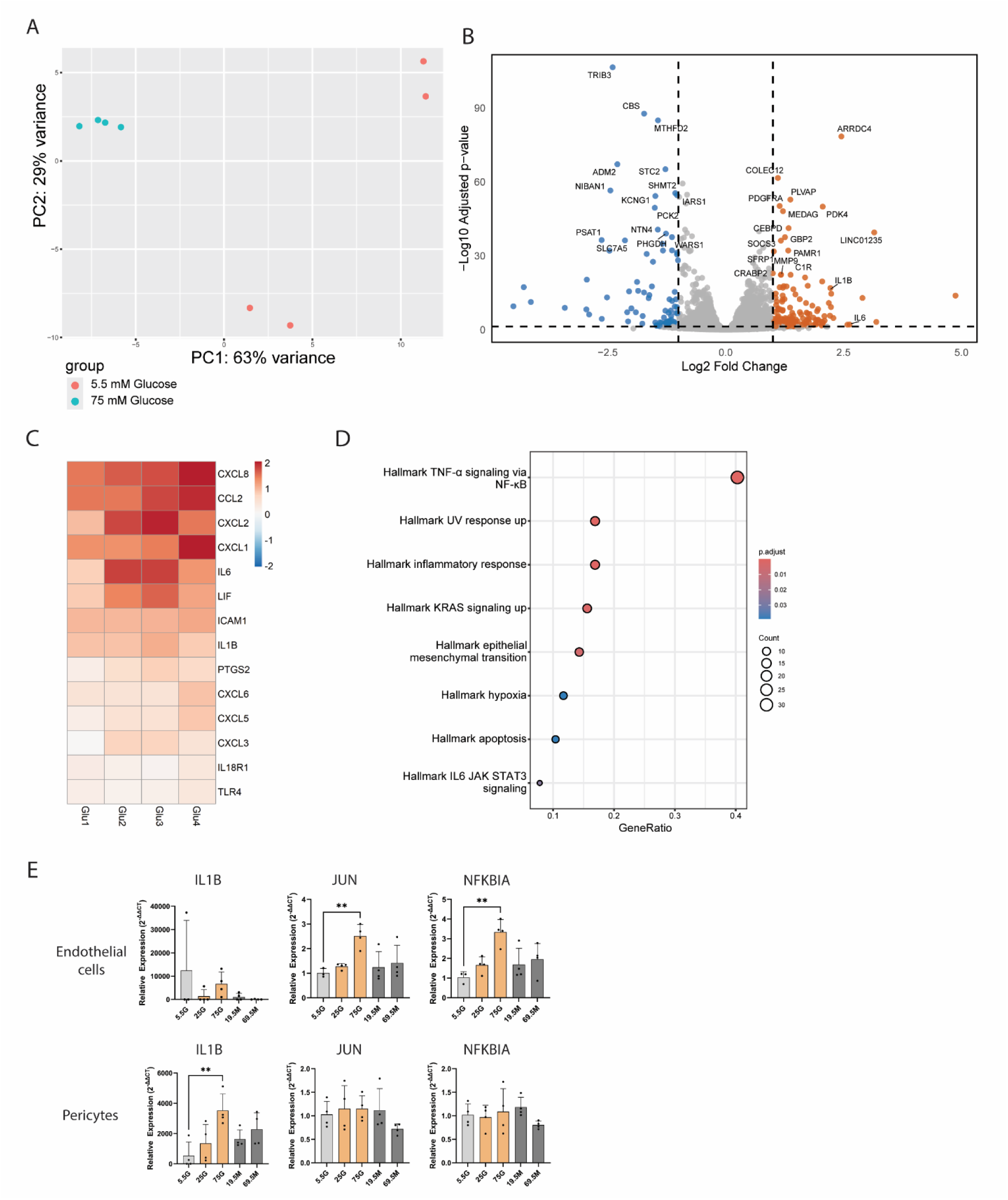
**Gene expression changes in CNS-mv-on-a-chip induced by high glucose level.** (A) The principal component analysis (PCA) plot shows clustering of samples cultured under physiological (5.5 mM glucose) and hyperglycemic (75 mM glucose) conditions. (B) Volcano plot illustrating differentially expressed genes between physiological and hyperglycemic CNS-mv-on-a-chip conditions. Orange and blue dots represent significantly upregulated and downregulated genes, respectively (FDR < 0.05, |FC| >2). (C) Heatmap showing upregulated genes associated with inflammation. Values are shown as fold changes relative to control group mean. (D) Over-representation analysis (ORA) plot of genes upregulated under hyperglycemia conditions. (E) Quantitative PCR analysis of IL1B, JUN and NFKBIA expression in ECs and PCs isolated from CNS-mv-on-a-chip cultured under physiological 5.5 mM glucose concentration (5.5G), hyperglycemic conditions 25 mM (25G) and 75 mM glucose (75G), or corresponding osmotic controls with mannitol (19.5M and 69.5M). Gene expression was normalized to GAPDH and HMBS as housekeeping genes and calculated using the ΔΔCt method. Data are presented as mean ± SD, n = 3-4.

To determine whether specific gene sets were enriched under diabetic conditions, we performed over-representation analysis (ORA) using the MSigDB Hallmark gene sets. The most significantly enriched pathway was TNF-α signaling via NF-κB, highlighting the central role of inflammatory activation (Fig. 3D). Other enriched signatures included general inflammation pathways, encompassing upregulation of cytokines, chemokines, and their receptors.

To confirm the TNF-α pathway activation and the primary responsive cell type, we isolated ECs and PCs from CNS-mv-on-a-chip using MACS sorting, followed by RNA extraction and qPCR. CNS-mv-on-a-chips were cultured under moderate (25 mM glucose) or severe (75 mM glucose) hyperglycemic conditions. Corresponding mannitol osmotic controls (19.5 and 69.5 mM mannitol) were included to rule out the effects of osmotic changes. We quantified expression of JUN and NFKBIA, two TNF-α-responsive genes upregulated in RNA-seq dataset, as well as IL1B as a pro-inflammatory marker.

The TNF-α signaling was activated only in ECs, but not PCs, indicated by the overexpression of JUN and NFKBIA expression. In contrast, IL1B gene expression was upregulated only in PCs. The osmotic controls did not show any changes (Fig. 3E). Although the lower hyperglycemic conditions demonstrated the same tendencies, the significant changes were induced only by severe hyperglycemia. These cell-type specific response suggest that ECs act as primary sensors of hyperglycemic stress through TNF-α/NF-κB activation, while PCs response by IL-1β secretion, potentially amplifying inflammation.

Having identified TNF-α/NF-κB activation as a key pathway responding to hyperglycemic stress, we next aimed to validate whether direct exposure to inflammatory cytokines could recapitulate the microvascular pathology caused by high glucose levels.

### Activation of TNF-α signaling results in similar effects on microvasculature parameters as hyperglycemic conditions

Gene expression analysis revealed that hyperglycemic conditions activated inflammatory signaling pathways (Fig. 3C, D, E). Notably, TNF-α signaling was strongly upregulated, along with the pro-inflammatory cytokines IL6 and IL1B, in the hyperglycemic CNS-mv-on-a-chip model.

To investigate whether these inflammatory pathways contribute to microvascular changes, we examined effects of pro-inflammatory cytokine IL-6, IL-1β, or TNF-α exposure on vascular morphology. Following treatment, we quantified vascular morphology parameters, including vessel diameter, vascular coverage area, number of branches per mm^2^, average branch length, connectivity ratio, and total vessel length per mm^2^.

IL-6 treatment had no significant effect on any of the assessed vascular parameters. In contrast, IL-1β exposure led to mild vascular regression, evidenced by reduced vessel diameter and vascular coverage area (Fig. 4A, B). The most pronounced morphological changes were observed with TNF-α treatment, which significantly impaired microvasculature integrity: vessel diameter, coverage area, number of branches per mm^2^, and connectivity ratio were all reduced, while average branch length increased, suggesting the regression of finer vessel structures. These results align with the transcriptomic data, which showed TNF-α signaling via NF-κB as the most enriched pathway under hyperglycemic conditions (Fig. 3D). Given that both TNF-α signaling and IL-1β were upregulated in the RNA-seq dataset and both activate the NF-κB pathway, we selected IL-1β/TNF-α co-treatment (IT) to represent the inflammatory component of diabetic conditions in subsequent experiments.

**Figure 4:**
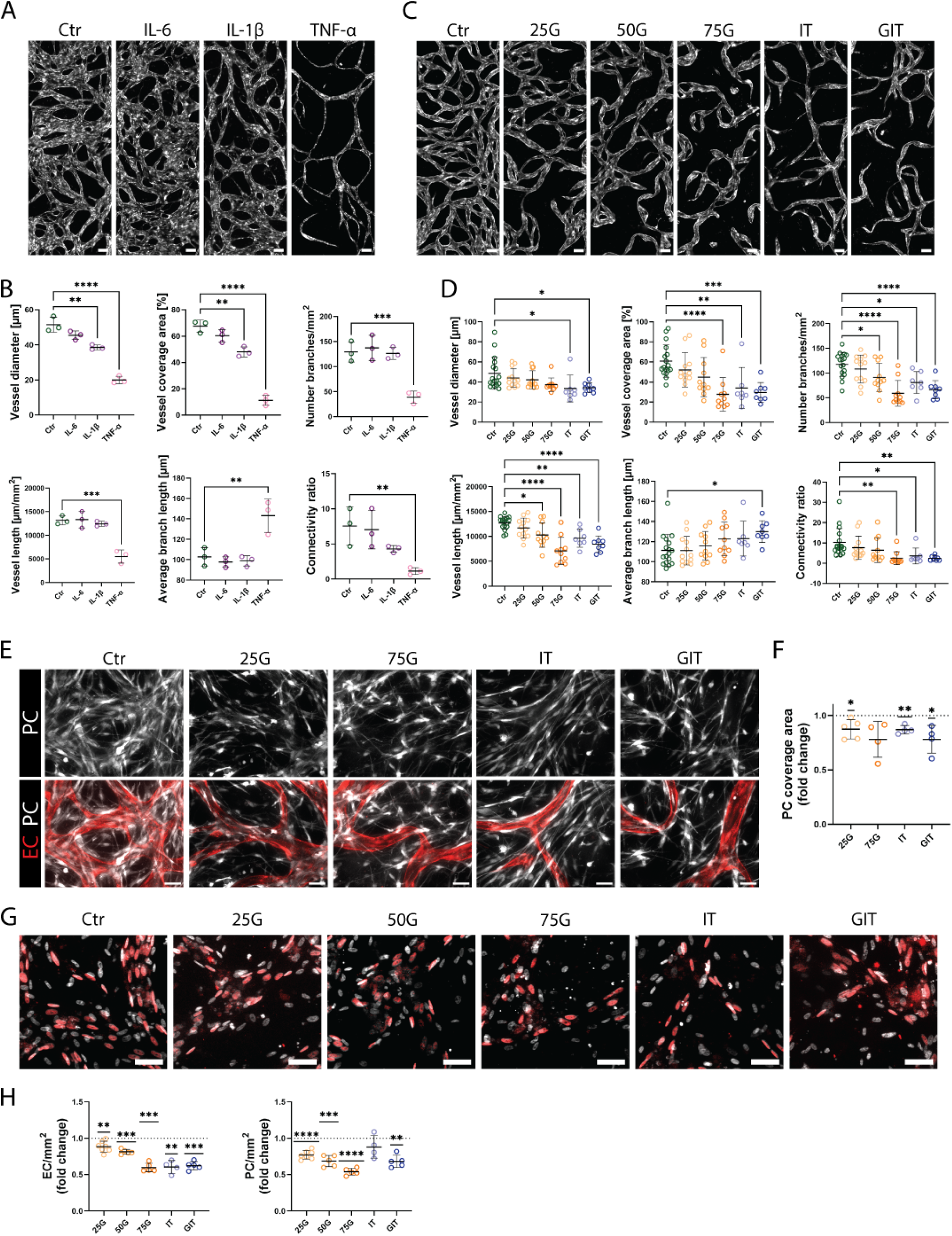
**Microvasculature morphology changes in CNS-mv-on-a-chip induced by diabetic conditions.** (A) Fluorescence images of CNS-mv-on-a-chip exposed to pro-inflammatory cytokines IL-6 (1 ng/ml), IL-1β (1 ng/ml), and TNF-α (1 ng/ml); ECs were labeled with mCherry (white). Scale bar: 100 µm. (B) Quantification of vascular morphology in CNS-mv-on-a-chip exposed to pro-inflammatory cytokines. N = 1, n = 3. (C) Fluorescence images of CNS-mv-on-a-chip under diabetic conditions: hyperglycemic treatments with 25 mM (25G), 50 mM (50G), and 75 mM (75G) glucose; treatment with inflammatory cytokines IL-1β (1 ng/ml) and TNF-α (1 ng/ml) (IT); combined diabetic condition with 25 mM glucose and IL-1β/TNF-α (GIT); control chips were cultured in medium with 5.5 mM glucose (5.5G). Scale bar: 100 µm. (D) Quantification of vascular morphology in CNS-mv-on-a-chip under diabetic conditions, N = 5-10, n = 7-12. (E) Fluorescence images of CNS-mv-on-a-chip used for PC coverage analysis; ECs were labeled with mCherry (red), and PCs with GFP (white). Scale bar: 50 µm. (F) Quantification of vascular area covered by PCs. Values are normalized to control group in each independent experiment. N = 4, n = 3-5. (G) Immunofluorescence images of CNS-mv-on-a-chip under diabetic conditions, stained with ERG-1/2/3 (red) to visualize EC nuclei, and Hoechst (white) for all nuclei. Scale bar: 100 µm. (H) Quantification of EC and PC cell numbers based on ERG-1/2/3 positive cells (ECs), and ERG-1/2/3 negative cells (PCs). Values were normalized to control chips cultured in medium with 5.5 mM glucose. N = 3-6, n = 4-7.

We next compared the effects of hyperglycemia and inflammation on microvasculature morphology. CNS-mv-on-a-chips were exposed to mild (25 mM glucose, 25G), moderate (50 mM glucose, 50G), or severe (75 mM glucose, 75G) hyperglycemic treatment. To rule out osmotic effects, osmotic controls with corresponding mannitol concentrations were included. The impact of hyperglycemia was compared with inflammatory background induced by IL-1β/TNF-α (IT). As a model of complete diabetic conditions, we applied 25 mM glucose with IL-1β/TNF-α (GIT).

CNS-mv-on-a-chips exposed to hyperglycemia and/or inflammation displayed clear signs of vascular regression (Fig. 4C). Both hyperglycemia and inflammatory conditions resulted in reduced vessel coverage area, number of branches per mm^2^, total vessel length, and connectivity ratio (Fig. 4D). The dose-dependent effect of hyperglycemia was observed: higher glucose concentrations caused more pronounced vascular alterations. Despite overall similarities, certain differences emerged – only inflammatory cytokines reduced vessel diameter, and only GIT treatment led to increased average branch length. Osmotic controls resulted in an occasional slight decrease in number of branches per mm^2^ and total vessel length (Supplementary Fig. 3A, B), but to a much lower degree than hyperglycemic conditions.

PC loss is one of the earliest hallmarks of microvascular regression. We aimed to determine whether signs of PC loss could be detected in the CNS-mv-on-a-chip model under diabetic conditions. As expected, hyperglycemic conditions (25G, 75G) and an inflammatory background (IT, GIT) reduced PC coverage of microvasculature compared to physiological controls (Fig. 4E, F). Mannitol-treated osmotic controls showed no changes in PC coverage (Supplementary Fig. 3C, D).

The observed reduction in PC coverage and vascular regression could reflect a decline in EC and/or PC numbers. To evaluate ECs number, EC nuclei were labeled using ERG-1/2/3 immunostaining. PC nuclei were identified as negative for ERG-1/2/3 staining. The number of ECs per mm² was significantly decreased under all diabetic conditions, including hyperglycemia (25G, 50G, and 75G), inflammatory background (IT), and combined diabetic condition (GIT) conditions (Fig. 4G, H). Perivascular PCs demonstrated similar tendencies, except that inflammatory conditions alone (IT) had a milder effect and did not significantly alter the number of PCs per mm^2^. EC and PC numbers were not altered by osmotic controls (Supplementary Fig. 3E, F).

Next, we focused on EC behavior in CNS-mv-on-a-chip under diabetic conditions. Vascular regression is characterized by the loss of ECs, while the basement membrane scaffold remains intact, forming “ghost vessels” – a non-perfused, non-functional capillary network. Under physiological conditions ghost vessels were rarely observed (Fig. 5A). Both hyperglycemic and inflammatory conditions induced ghost vessel formation (Fig. 5A) and significantly increased the ghost vessel fraction under inflammatory condition (IT) (Fig. 5B). Osmotic controls did not significantly alter the ghost vessel fraction (Supplementary Fig. 4A, B).

**Figure 5:**
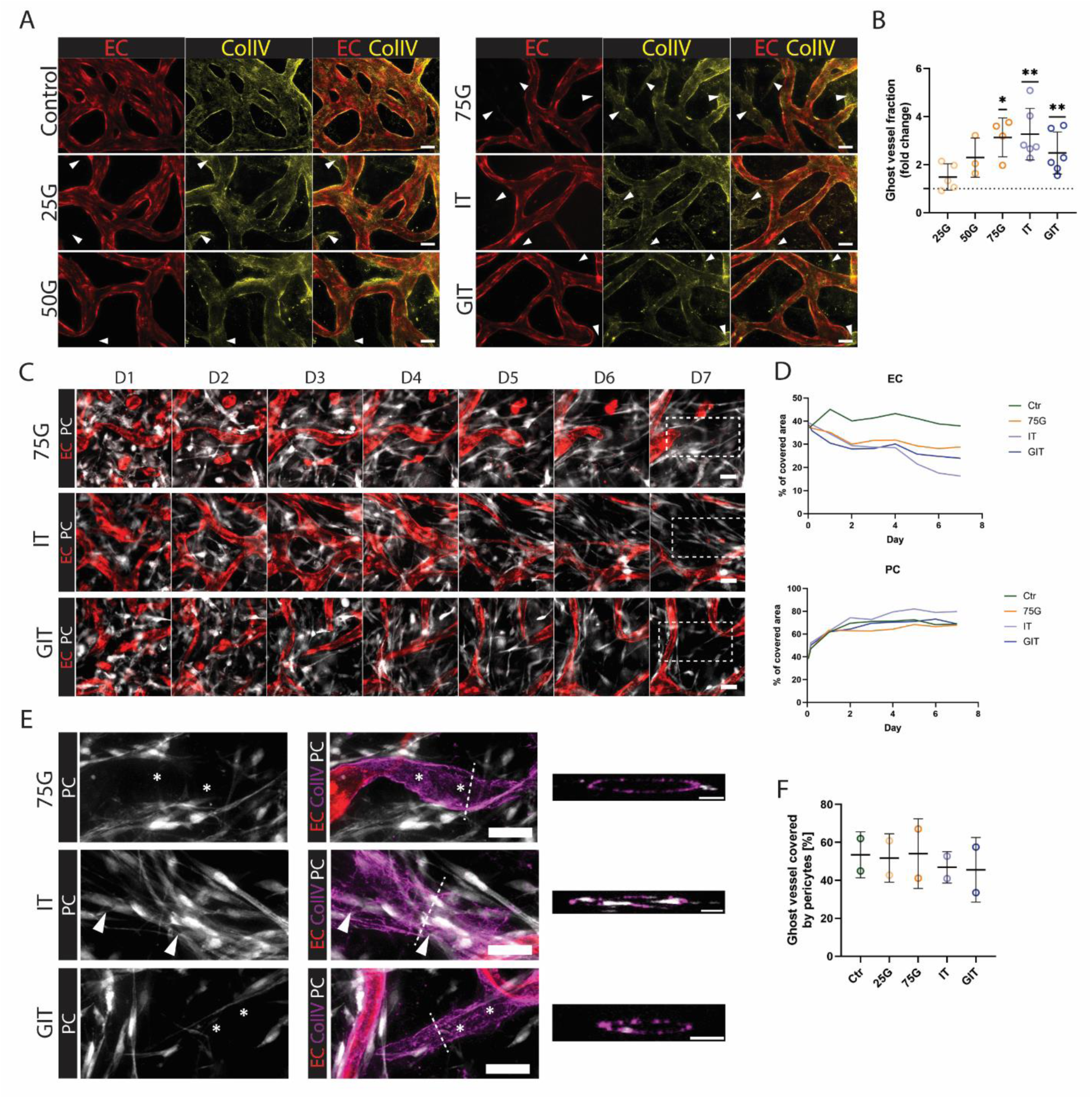
**Diabetic conditions induced ghost vessels.** (A) Representative images of ghost vessels in CNS-mv-on-a-chip under diabetic conditions, including hyperglycemic treatments with 25 mM (25G), 50 mM (50G), and 75 mM (75G) glucose; treatment with inflammatory cytokines IL-1β /TNF-α (1 ng/ml) (IT); combined diabetic condition with 25 mM glucose and IL-1β/TNF-α (GIT); control chips were cultured in medium with 5.5 mM glucose. ECs were labeled with mCherry (red); basement membranes were visualized via immunostaining for collagen IV (yellow). White arrows indicate ghost vessels. Scale bar: 50 µm. (B) Quantification of the ghost vessel fraction under diabetic conditions. N = 3-6, n = 3-6. (C) Representative images of regressing vessels under diabetic conditions over 7 days. ECs were labeled with mCherry (white), and PCs were labeled with GFP (white). Dashed line marked the region shown on Fig. 5E. Scale bar 50 µm. (D) Quantification of area fraction covered by ECs and PC over 7 days under diabetic conditions. (E) Immunostaining showing ghost vessels. ECs were labeled with mCherry (red), and PCs were labeled with GFP (white), basement membranes were visualized via immunostaining for collagen IV (magenta). The dashed line marks the cross-section, the orthogonal view of the z-stack across this dashed line is shown on the right. Asterisks mark ghost vessels not covered by PCs. Scale bar 50 µm. (F) Quantification of the percentage of ghost vessels fraction, covered by PCs. N = 1, n = 1-2.

**Figure 6:**
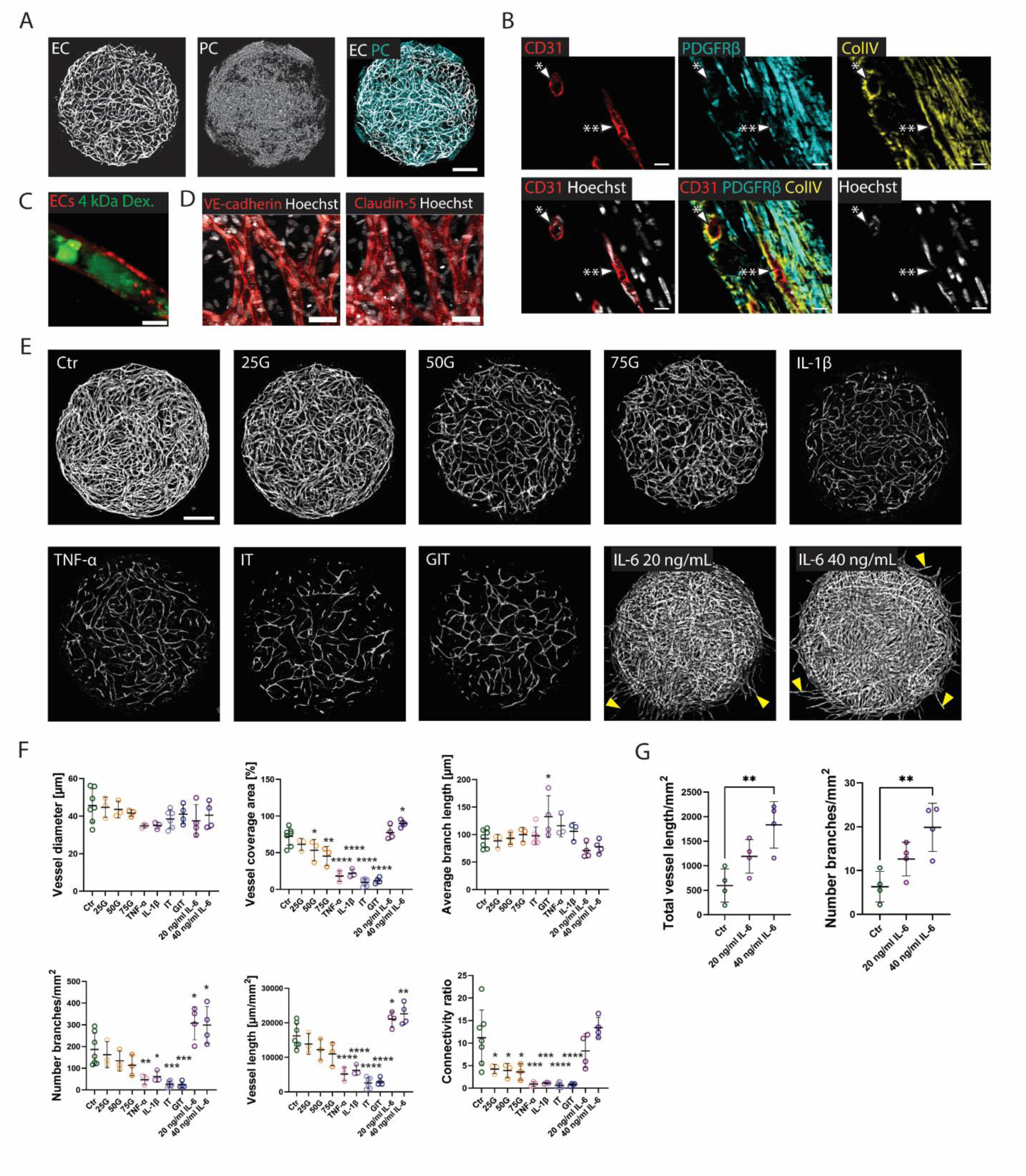
**Effects of diabetic conditions on microvascular drops.** (A) Fluorescence images showing microvascular drop model. ECs were labeled with mCherry (white), PCs were labeled with GFP (white or cyan). Scale bar: 500 µm. (B) Immunostaining of cryosectioned microvascular drop showing EC marker CD31 (red), PC marker PDGFRβ (cyan), basement membrane marker ColIV (yellow). * indicates a transverse and ** a longitudinal section through vessels. Scale bar: 20 µm. (C) Fluorescence image showing the microvascular lumen perfused with with 4 kDa FITC-dextran (green). ECs were labeled with mCherry (red). Scale bar: 10 µm.(D) Immunostaining of microvasculature on drop showing adherens junction marker VE-cadherin and tight junction marker claudin-5 (red). Scale bar: 50 µm. (E) Fluorescent images of microvascular drops after 7 days of exposure to 25 mM (25G), 50 mM (50G), and 75 mM (75G) glucose; IL-β, 1, TNF-α, IL-β/TNF-α (IT), combined diabetic conditions (GIT); or 48-hour exposure to 20 and 40 ng/ml IL-6. Control drops were cultured in 5.5 mM glucose. ECs were labeled with mCherry (white). Yellow errors indicate sprouting vessels. Scale bar: 500 µm. (F) Quantification of microvascular morphology in microvascular drops. N = 1-6, n = 3-9 (G) Quantification of sprouting vessels length. N = 1, n = 4.

To visualize the dynamics of vessel regression, ECs were labeled with mCherry, PCs with GFP, and the total cellular area of CNS-mv-on-a-chips was imaged over 7 days (Fig. 5C). Both severe hyperglycemia (75G) and inflammatory conditions (IT and GIT) induced a visible decrease in the total EC-covered area (Fig. 5D). The area covered by PCs remained unchanged, suggesting that the reduction in PC coverage observed during inflammation is more likely due to impaired PC recruitment or attachment, rather than reduction in PC number alone. Area covered by ECs and PCs were unchanged in osmotic control conditions (Supplementary Fig. 4C). We additionally confirmed that roughly half of the ghost vessel area remained scaffolded by PCs (Fig. 5E, F) and diabetic conditions did not change the fraction of ghost vessels covered with PCs (Fig. 5F). Osmotic control conditions did not alter ghost vessel coverage by PCs (Supplementary Fig. 4D)

In conclusion, both hyperglycemia and inflammatory conditions induced vascular regression, decrease PC coverage of vascular network, and decreased both EC and PC cell numbers. However, based on the vessel regression dynamics, the regression appears to be primary driven by EC retraction, while PCs might partly remain attached and scaffold ghost vessels after EC regression.

### Microvascular drops as a complementary platform for analyzing vascular morphology

Our results demonstrate that the CNS-mv-on-a-chip provides an advanced model for studying microvasculature under diabetic conditions. However, large-scale screenings might require cost-effective solutions to reduce both the time and costs of drug testing. Therefore, we developed a complementary three-dimensional microvascular drop system compatible with 96-well plate format, and compared the effects of diabetic conditions in the microfluidic chip and microvascular drops to evaluate their reproducibility across platforms and accuracy.

To form microvascular drops, ECs and PCs were incorporated into a fibrin matrix. Microvascular drops demonstrated a three-dimensional capillary network (Fig. 7A), which also contained a defined lumen (Fig. 7C). Cryosectioning of the drops confirmed that the capillaries not only contained a lumen but were also surrounded by PCs (PDGFRβ staining) and formed a basement membrane (Collagen IV staining) (Fig. 7B). The capillary network was positively stained for the EC marker CD31, the adherent’s junction marker VE-cadherin, and tight junction marker claudin-5 (Fig. 7D).

We next tested how diabetic conditions influence microvascular morphology, and whether these results are comparable with those from the CNS-mv-on-a-chip. The microvasculatures were exposed to mild 25 mM glucose (25G), moderate 50 mM glucose, and severe 75 mM glucose (75G) hyperglycemic treatment. The inflammatory background was induced by inflammatory cytokines IL-1β and/or TNF-α (IT), and combined diabetic conditions, which included 25 mM glucose and IL-1β/TNF-α (GIT). Similar to the CNS-mv-on-a-chip, both hyperglycemia and inflammatory conditions caused signs of vascular regression (Fig. 7E, F). All hyperglycemic conditions (25G, 50G, and 75G) reduced the vasculature connectivity ratio, while moderate and severe hyperglycemia (50G, and 75G) also reduced the vessel coverage area. All inflammatory conditions (TNF-α, IL-1β, IT, and GIT) reduced the number of branches per mm^2^ and vessel length. Interestingly, in line with results obtained from the CNS-mv-on-a-chip, only GIT treatment led to an increased average branch length (Fig. 4D, Fig. 7F). In addition, we did not observe any significant changes in diameter, however, the observed trends were consistent with the CNS-mv-on-a-chip results: inflammatory background had much more pronounced effect than hyperglycemia. The mannitol osmotic control did not show any significant changes (Supplementary Fig. A, B).

We then tested the effect of IL-6 on microvasculature formation. The CNS-mv-on-a-chip did not show signs of vascular regression after seven days of treatment with 1 ng/ml IL-6 (Fig. 4A). Therefore, we aimed to test whether acute, short-term (48 h) treatment with higher concentrations (20 ng/ml and 40 ng/ml) would have any effects on the microvasculature.

Interestingly, increased IL-6 concentrations had an opposite effect on the microvasculature compared to TNF-α and/or IL-1β and hyperglycemia: IL-6-treated drops demonstrated increased total vessel length and number of branches per mm² (Fig. 7E, F). Moreover, we observed enhanced vessel sprouting from the microvascular drops upon treatment (Fig. 7E, yellow arrows, G).

In summary, we successfully replicated the vascular morphological changes observed under hyperglycemia and inflammatory conditions in the CNS-mv-on-a-chip using microvasculature drop model, confirming high reproducibility across two platforms. Moreover, the microvasculature on a drop provided an additional opportunity to measure vessel sprouting, as the vasculature network was not enclosed in a chip geometry.

### Introduction of macroglia in the CNS-mv-on-a-chip model

The CNS-mv-on-a-chip model successfully replicated the hallmarks of diabetes-associated microvascular pathologies, such as PC loss, vascular regression, and ghost vessel formation. We then aimed to further improve the model, by incorporating macroglial cells, which are known to support microvasculature architecture in the brain and retina. We received and characterized hiPSC-derived astrocytes (ACs), and Müller glia cells (MGs). The identity of ACs were confirmed by star-shape morphology and expression of markers, including GFAP, S100β, AQP4, and CD44. (Supplementary Fig. 6B) MGs were characterized by the presence of CD29, CD44, Dkk3, and glutamine synthetase (Supplementary Fig. 6A).

Both hiPSC-derived ACs and MGs successfully integrated into the CNS-mv-on-a-chip (Fig. 8A, B). MGs (Fig. 8A) and ACs (Fig. 8B) were observed in close contact with ECs and PCs, wrapping their endfeet around vasculature (white arrows). This close contact between glia cells and vasculature suggests the cells successfully structural integrations.

**Figure 7:**
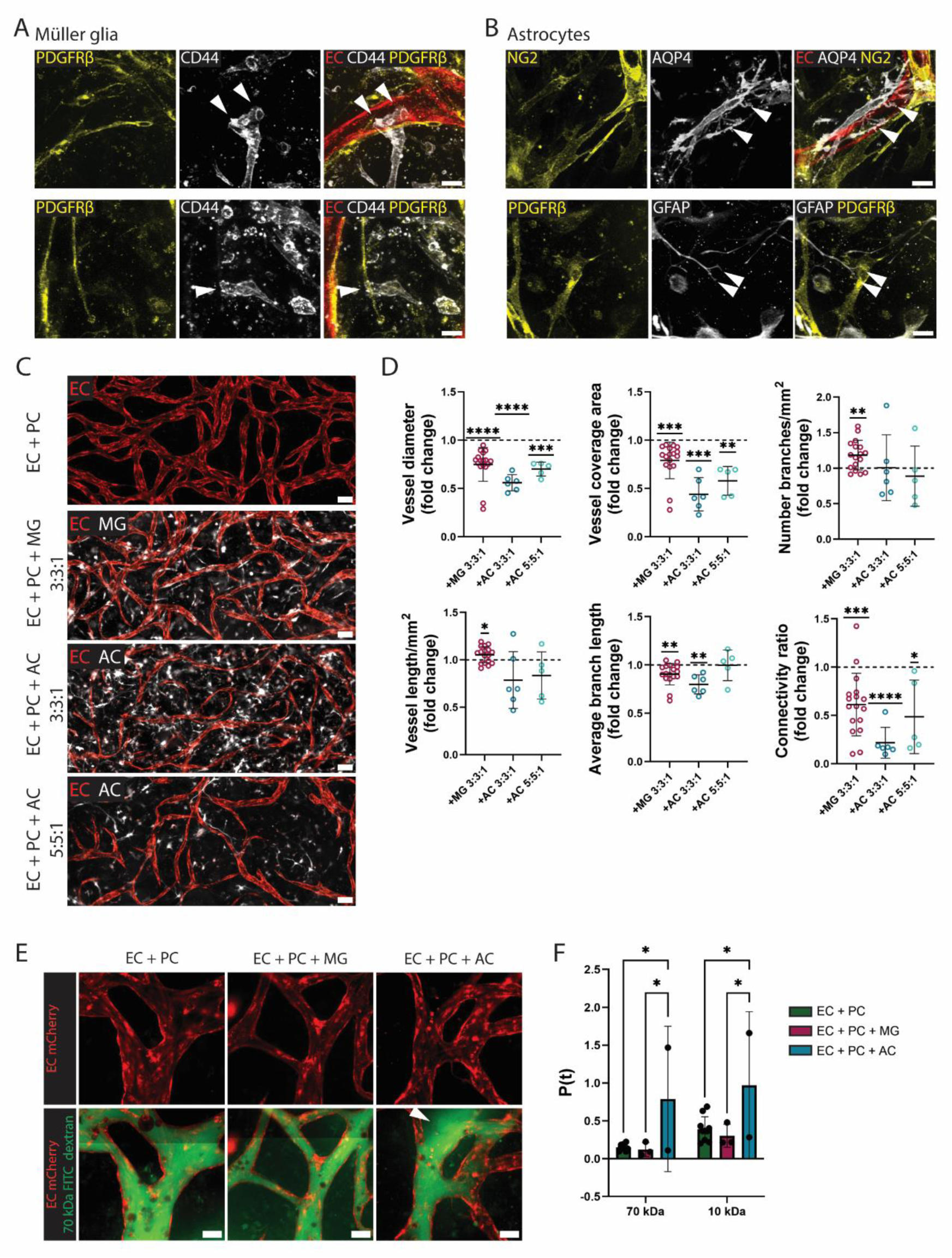
**Introduction of macroglial cells into the CNS-mv-on-a-chip.** (A) Immunostaining of CNS-mv-on-a-chip formed using ECs, PCs, and Müller glial cells (MGs). ECs were labeled with mCherry (red); PCs were stained with PDGFRβ marker (yellow), and MGs with CD44 (white). Scale bar: 20 µm. White errors mark contacts of endfeet with vasculature. (B) Immunostaining of CNS-mv-on-a-chip formed using ECs, PCs, and astrocytes (ACs): ECs were labeled with mCherry (red), PCs were stained with PDGFRβ marker (yellow), ACs with GFAP (white). White arrows mark contacts of glial endfeet with the vasculature. Scale bar: 20 µm. (C) Immunostaining showing AC morphology in CNS-mv-on-a-chip. ECs were labeled with mCherry (red), ACs were stained for GFAP (white) Scale bar: 50 µm. (D) Fluorescence images of CNS-mv-on-a-chip with glia incorporation. ECs were labeled with mCherry (red), and ACs or MGs with GFP (white). Scale bar: 100 µm. (E) Quantification of vascular morphology parameters in CNS-mv-on-a-chip with integrated glial cells. N = 3-6, n = 3-20. (F). Fluorescence images of the CNS-mv-on-a-chip perfused with 70 kDa FITC-dextran (green). ECs were labeled with mCherry (red). Scale bar: 50 µm. (G) Quantification of the apparent permeability coefficient. N = 1-3, n = 2-10.

**Figure 8:**
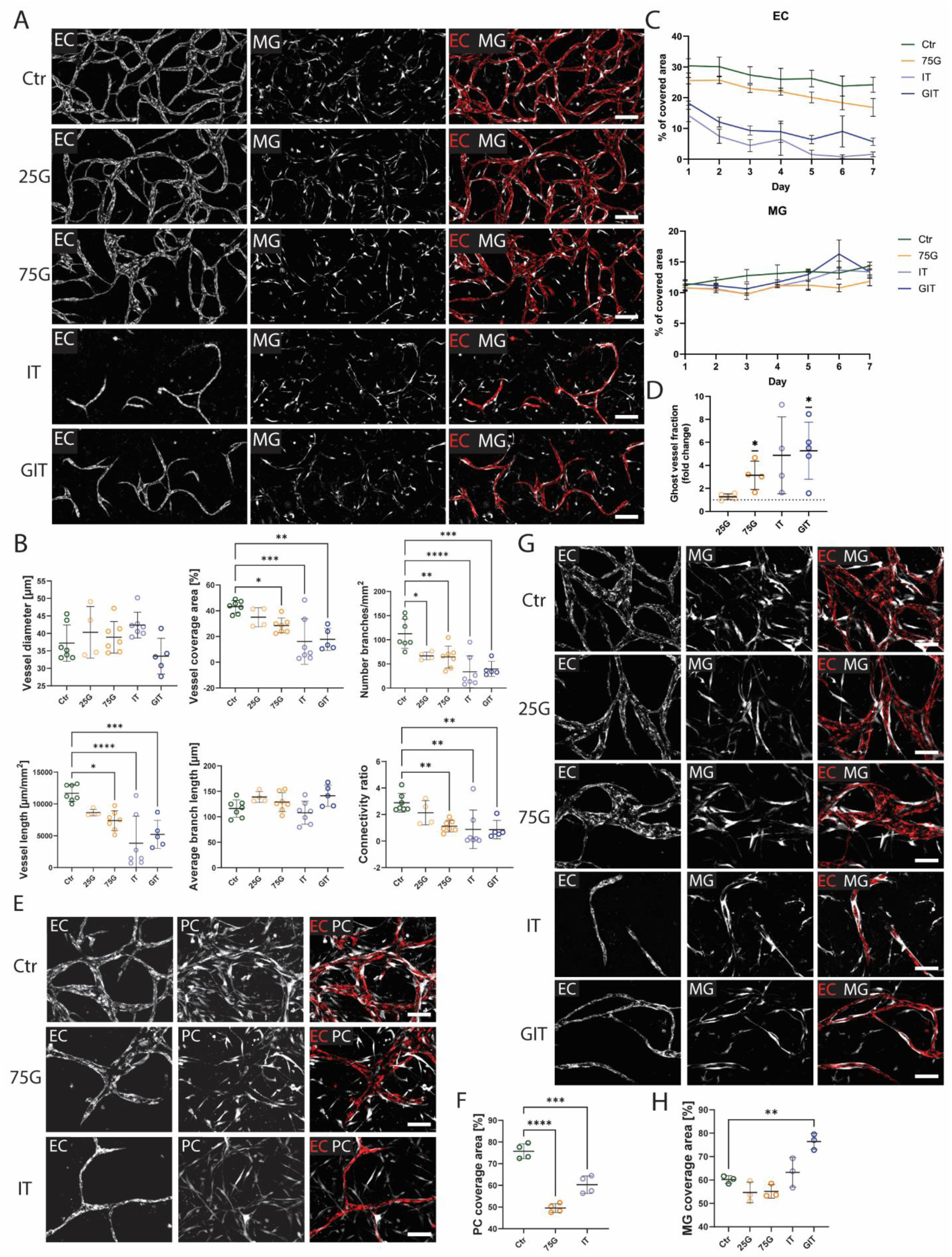
**Morphological changes in the inner blood-retinal barrier (iBRB)-mv-on-a-chip with Müller glia (MG) under diabetic conditions.** (A) Fluorescence images of CNS-mv-on-a-chip with integrated MGs under diabetic conditions: hyperglycemic treatments with 25 mM (25G), 50 mM (50G), and 75 mM (75G) glucose; treatment with inflammatory cytokines IL-1β (1 ng/ml) and TNF-α (1 ng/ml) (IT); combined diabetic condition with 25 mM glucose and IL-1β/TNF-α (GIT). Control chips were cultured in medium with 5.5 mM glucose (5.5G). ECs were labeled with mCherry (red) and MGs with GFP (white). Scale bar: 200 µm. (B) Quantification of vascular morphology in iBRB-mv-on-a-chip under diabetic conditions. N = 4, n = 4-7. (C) Quantification of the area fraction covered by ECs and MGs over 7 days under diabetic conditions. N = 1, n = 3. (D) Quantification of the ghost vessel fraction under diabetic conditions. N = 3, n = 4-5. (E) Fluorescence images of iBRB-mv-on-a-chip used for MG coverage analysis. ECs were labeled with mCherry (red) and MGs with GFP (white). Scale bar: 100 µm. (F) Quantification of the vascular area covered by MGs. N = 1, n = 3. (G) Fluorescence images of iBRB-mv-on-a-chip used for PC coverage analysis. ECs were labeled with mCherry (red) and PCs with GFP (white). Scale bar: 100 µm. (H) Quantification of the vascular area covered by PCs. N = 1, n = 3.

Next, we analyzed the influence of incorporated glial cells on the vascular morphology of CNS-mv-on-a-chip. The physiological 1:1 ratio of ECs to PCs in CNS vasculature was maintained in the CNS-mv-on-a-chip while adding glial cells, resulting in a final EC:PC:AC/MG ratio of 3:3:1. Glial incorporation significantly decreased vessel diameter, vessel coverage area, and connectivity (Fig. 8C, D). Interestingly, ACs demonstrated stronger effects on vascular parameters. To be able to compare the effects of MGs and ACs, we reduced the amount of ACs to the 5:5:1 ratio, which produced effects more comparable to MGs (Fig. 8C, D), and this ratio was used for further functional comparison. Additionally, only MGs increased the total vessel length and number of branches per mm^2^, indicating that MGs in comparison to ACs may improve the complexity of microvasculature network.

Since we observed a successful structural integration of glial cells into the CNS-mv-on-a-chip, we next assessed the barrier function in CNS-mv-on-a-chip using fluorescently labeled 70 kDa FITC-dextran and 10 kDa AF-640-dextran (Fig. 8E). MG integration did not change the apparent permeability coefficient (Fig. 8F) (ECs+PCs: 0.15 ± 0.04 µm/s for 70 kDa dextran and 0.39 ± 0.16 µm/s for 10 kDa dextran; ECs+PCs+MG: 0.12 ± 0.09 µm/s for 70 kDa dextran and 0.30 ± 0.15 µm/s for 10 kDa dextran), suggesting that the CNS-mv-on-a-chip formed by ECs and PCs was already tight, and the addition of MGs did not improve barrier function. In contrast, ACs significantly increased the apparent permeability coefficient (0.79 ± 0.96 µm/s for 70 kDa dextran and 0.97 ± 0.97 µm/s for 10 kDa dextran). This unexpected effect of AC incorporation likely reflects an immature or reactive state of the hiPSC-derived ACs. This result is consistent with previous findings showing that hiPSC-derived ACs often exhibit an activated, pro-inflammatory phenotype, which can disrupt microvascular barrier function [24].

In summary, hiPSC-derived MGs were successfully integrated into the CNS-mv-on-a-chip, resulting in decreased vessel diameter, coverage area, and connectivity, increasing total vessel length and number of branches per mm^2^, while preserving the barrier function. hiPSC-derived ACs, despite structural integration, disrupted CNS-mv-on-a-chip barrier function, that had been already shown in previous studies [24].

### Analysis of inner blood retinal barrier (iBRB)-mv-on-a-chip with Müller glia cells under diabetic conditions

Given that MGs modulated vascular morphology without compromising barrier function, we selected MGs as a macroglial component to advance our CNS microvascular model towards a retinal identity and replicated the inner blood-retinal barrier (iBRB) composition. The iBRB-mv-on-a-chip was exposed to hyperglycemic conditions (25 mM, 50 mM, and 75 mM glucose), inflammatory conditions induced by IL-1β and TNF-α co-treatment (IT), and combined diabetic conditions with 25 mM glucose and TNF-α/IL-1β (GIT). The iBRB-mv-on-a-chip responded to diabetic conditions with clear signs of vascular regression (Fig. 9A). Vascular morphology analysis confirmed vascular regression of the iBRB-mv-on-a-chip under diabetic conditions: both hyperglycemia and inflammatory conditions reduced vessel coverage area, number of branches per mm², total vessel length, and connectivity ratio (Fig. 9B). In contrast to the CNS-mv-on-a-chip, diabetic conditions did not alter blood vessel diameter in the iBRB-mv-on-a-chip (Fig. 4D), most likely because the vessel diameter was already smaller due to MG integration (Fig. 8D). Mannitol osmotic controls showed no significant changes in vascular morphology (Supplementary Fig. 7A, B), confirming that the observed effects were not caused by osmotic stress.

Vessel regression dynamics demonstrated that both severe hyperglycemia (75G) and inflammatory conditions (IT and GIT) induced a visible decrease in the EC-covered area (Fig. 9C), with the inflammatory background causing a more profound negative effect. The microvasculature area covered with MGs remained unchanged under all conditions, suggesting that diabetic conditions did not induce MG cell death. Osmotic control conditions with mannitol did not cause a change in area covered by ECs or MGs (Supplementary Fig. 7C). Moreover, severe hyperglycemia (75G) and inflammation (IT and GIT) increased the ghost vessel fraction (Fig. 9D), further confirming vascular regression induced by diabetic conditions. Osmotic control condition with 69.5 mM mannitol resulted in increased ghost vessel fraction, however to a much smaller degree then hyperglycemic conditions (Supplementary Fig. 7D).

Since we observed that diabetic conditions reduced PC coverage in the CNS-mv-on-a-chip (Fig. 4E, F), we quantified PC coverage in the iBRB-mv-on-a-chip. Incorporation of MGs did not change PC coverage of iBRB microvasculature (76.1 ± 6.5 %) compared with CNS microvasculature formed by ECs and PCs (75.5 ± 4.0 %). In line with results obtained from the CNS-mv-on-a-chip, both hyperglycemia and inflammation significantly reduced PC coverage in the iBRB-mv-on-a-chip (Fig. 9E, F).

In the iBRB, MGs formed a sheath around capillaries. To evaluate the effect of diabetic conditions on MG coverage, we quantified the capillary network fraction covered by MGs. Interestingly, hyperglycemia did not change MG coverage of vessels, while inflammation (IT), and even more strongly combined diabetic induction (GIT), enhanced the MG sheath around capillaries (Fig. 9G, H). MG coverage was not influenced by osmotic control conditions (Supplementary Fig. 7E, F).

The obtained results demonstrate that MGs did not exert protective or harmful effects on the microvasculature under diabetic conditions, as microvascular morphology changes were comparable to both CNS-mv-on-a-chip and microvasculature drop system. Although diabetic vessel regression in iBRB-mv-on-a-chip was accompanied by PC coverage reduction, MGs remained in close contact with capillaries and even enhanced their sheath around vessels, suggesting altered MG behavior under diabetic conditions.

## Discussion

The microvasculature is a crucial part of the cardiovascular system, consisting of small arterioles, venules, and capillary networks. Capillaries play an essential role in delivering nutrients, oxygen, and signaling molecules to tissues, while restricting potentially dangerous substances. This protective barrier is particularly tight in the central nervous system (CNS). During aging and in the course of diseases, such as small vessel disease and diabetic retinopathy, the CNS microvasculature (CNS-mv) is affected: these vessels are characterized by a reduced number of perivascular pericytes (PC), vascular regression, and leaky capillaries [38–40]. This dysfunction leads to severe consequences, such as strokes or blindness. Current *in vitro* microvasculature models struggle to capture the multicellular interactions and dynamic processes underlying microvascular disease. Here, we address this gap by developing three complementary *in vitro* CNS-mv models that recapitulate key features of diabetic microvascular pathology and can be used to investigate therapeutic targets.

Our *in vitro* CNS-mv models demonstrated high reproducibility of microvascular pathologies caused by diabetic conditions across platforms (Table 1) and were designed to complement each other in the process of drug development. The drop model allows cost-effective and rapid evaluation of microvasculature morphology, enabling large-scale screening. The CNS-mv-on-a-chip provides a platform for detailed microvasculature analysis, including endothelial cell (EC) to pericyte (PC) ratios, PC coverage, ghost vessel formation, and vascular regression dynamics, thereby supporting mechanistic studies. The inner blood-retinal barrier (iBRB)-on-a-chip model enables retina-specific investigations and the testing of therapies targeting Müller glial cells. Together, these platforms model key aspects of human microvasculature and offer versatile tools for future research.

**Table 1.** Comparative effects of diabetic conditions across three microvascular in vitro models.

|  | Model | CNS-mv-on-a-chip | Microvascular Drops | iBRB-mv-on-a-chip |
| --- | --- | --- | --- | --- |
| Parameter | Condition |  |  |  |
| Vessel Diameter | Hyperglycemia (25-75G) | No change | No change | No change |
|  | Inflammation (IT) | Decreased | No change | No change |
|  | Combined (GIT) | Decreased | No change | No change |
| Vessel Coverage Area | Hyperglycemia (25-75G) | Decreased | Decreased | Decreased |
|  | Inflammation (IT) | Decreased | Decreased | Decreased |
|  | Combined (GIT) | Decreased | Decreased | Decreased |
| Number of Branches | Hyperglycemia (25-75G) | Decreased | No change | Decreased |
|  | Inflammation (IT) | Decreased | Decreased | Decreased |
|  | Combined (GIT) | Decreased | Decreased | Decreased |
| Total Vessel Length | Hyperglycemia (25-75G) | Decreased | No change | Decreased |
|  | Inflammation (IT) | Decreased | Decreased | Decreased |
|  | Combined (GIT) | Decreased | Decreased | Decreased |
| Average Branch Length | Hyperglycemia (25-75G) | No change | No change | No change |
|  | Inflammation (IT) | No change | No change | No change |
|  | Combined (GIT) | Increased | Increased | No change |
| Connectivity Ratio | Hyperglycemia (25-75G) | Decreased | Decreased | Decreased |
|  | Inflammation (IT) | Decreased | Decreased | Decreased |
|  | Combined (GIT) | Decreased | Decreased | Decreased |
| Ghost Vessel Fraction | Hyperglycemia (25-75G) | Increased | Not measurable | Increased |
|  | Inflammation (IT) | Increased | Not measurable | Increased |
|  | Combined (GIT) | Increased | Not measurable | Increased |
| PC coverage | Hyperglycemia (25-75G) | Decreased | Not measurable | Decreased |
|  | Inflammation (IT) | Decreased | Not measurable | Decreased |
| MG coverage | Hyperglycemia (25-75G) | Not measurable | Not measurable | No change |
|  | Inflammation (IT) | Not measurable | Not measurable | Increased |

Although the main components of CNS capillaries – ECs and PCs – can be obtained from human donors, their scarcity and some degree of variability between donors make research work based on primary cells difficult. In particular, CNS PCs are hard to obtain, since PCs of the CNS microvasculature have a different origin from PCs of the rest of the body [41] and therefore have to be isolated from brain tissue. To overcome the difficulties of using donor cells, we used hiPSC-derived cells.

We successfully established a physiologically relevant CNS-mv-on-a-chip model using hiPSC-derived ECs and neural crest-originated PCs. The CNS-mv-on-a-chip model recapitulated essential features of CNS-mv, such as physiological PC coverage (76.1 ± 6.5 %) [42] (Fig. 1B, E), tight junctions (Fig. 2A), basement membrane formation (Fig. 1E), and low permeability coefficients (Fig. 2C, D). In our system, we confirmed that PCs play a critical role in microvasculature formation: although ECs in a 3D gel were able to form a network of capillary-like structures, the EC-only microvasculature demonstrated a 10-fold increase in the permeability coefficients (Fig. 2D), highlighting that the PC coverage plays a critical role in supporting the morphology and function of the capillary network.

We used the CNS-mv-on-a-chip to model pathological changes under diabetic conditions. Transcriptomic analysis revealed inflammatory pathways activation, in particular TNF-α signaling via NF-κB (Fig. 3B, C, D). This finding is consistent with previously published data, showing NF-κB role in the progression of diabetes-associated microvascular pathologies [43–45]. Overrepresentation analysis identified enrichment of inflammatory response hallmark, and upregulation of pro-inflammatory genes, including CXCL8, CCL2, CXCL2, CXCL1, IL6, and IL1B (Fig. 3B, C). Such inflammatory activation is one of pathology-driving factors of diabetic vascular complications, including macular edema, ischemia and neovascularization [46, 47]. Similar results of TNF-α signaling activation under diabetic conditions has been already received using human primary retinal cells in the chip [23], confirming that hiPSC-derived cells can be successfully used to replicate human donor cell DR phenotypes.

Our cell-type-specific qPCR analysis revealed inflammatory crosstalk: TNF-a/NF-κB signaling was activated in ECs, whereas PCs showed elevated IL-1β expression (Fig. 3E). Elevated IL-1β level has been reported in vitreous fluid of patients with diabetes [48] and in diabetic rat model [49]. *In vitro*, PCs released IL-1β in response to ischemic injury stress [50]. Furthermore, PCs express IL-1 receptors and respond to IL-1β stimulation by producing inflammatory mediators, including TNF-α [51], IL-6 [50, 52], and NF-κB [53], resulting in a positive feedback loop that amplifies neuroinflammation. Our data suggest that hyperglycemia primary affects ECs *via* TNF-α/NF-κB pathway activation, while PCs potentially amplify the inflammatory pathway through IL-1β release.

In line with these findings, direct TNF-α exposure induced severe vascular degeneration (Fig. 4 A), marked by reduced vessel diameter, decreased vessel coverage area, total vessel length, number of branches per mm^2^, and connectivity (Fig. 4B). IL-1β treatment induced similar, but milder vascular damage. Together, these findings identify a PC-EC inflammatory axis, mediated by IL-1β and TNF-α/ NF-κB, as a potential driver of diabetic microvascular pathology and suggest that blocking IL-1β/ TNF-α signaling as promising therapeutic targets to prevent microvasculature pathologies in CNS. The therapeutic application of these findings may be substantial, as IL-1β (Canakinumab, Rilonacept) and TNF-α (Infliximab, Adalimumab, Etanercept, Golimumab, Certolizumab) antagonists are already FDA-approved for other indications. Indeed, anti-TNF-α therapy has already been tested for macular edema [54]. Our results suggest TNF-α and IL-1β as targets for preventing DR pathologies under hyperglycemic conditions.

Subsequently, we used co-treatment with IL-1β and TNF-α (IT) to model the inflammatory component of diabetes, and combined hyperglycemia (25 mM glucose) and inflammation to model full diabetic conditions (GTI). In CNS-mv-on-a-chip, both hyperglycemia and inflammation induced similar microvascular degeneration (Fig. 4C): both conditions reduced vessel coverage, number of branches per mm^2^, total vessel length, and connectivity (Fig. 4D). However, some differences emerged: only inflammatory conditions (IT and GIT) reduced vessel diameter, whereas only the combined diabetic condition (GIT) increased average branch length.

PC coverage was reduced under both hyperglycemic and inflammatory conditions (Fig. 4E, F). Both EC and perivascular PC cell numbers were decreased in diabetic conditions in the CNS-mv-on-a-chip (Fig. 4G, H), while total cell number was only impacted in case of ECs (Fig. 5 D). These results suggest that inflammatory cytokines impair PC recruitment or EC-PC crosstalk rather than drastic decrease in PC number. This interpretation aligns with published data, showing that TNF-α promotes the migration and remodeling of cerebral PCs [55].

Retraction of ECs, leaving the empty basement membrane scaffold – so-called ghost vessels – is a hallmark of CNS vascular pathology [56, 57]. Although ghost vessels are well documented in postmortem tissues, the dynamics of their formation remain unclear due to limitations in real-time observation. Consistent with *in vivo* pathology, we observed the ghost vessel formation under both severe hyperglycemia (75G) and inflammation (IT) (Fig. 5A). Life imaging of CNS-mv-on-a-chip confirmed that vascular regression was initiated by EC retraction, while PCs partially remained associated with ghost vessels (Fig. 5C, D). These observations provide an insight into the ghost vessel formation dynamics: hyperglycemia and/or inflammation disrupt EC-PC contacts, marked by reduced PC-coverage; this triggers EC retraction and ghost vessel formation, with some PCs remaining temporarily attached to the basement membrane; ultimately, PC numbers decline, and ghost vessels lose PC coverage. The prolonged PC coverage during vessel regression suggest a therapeutic window, when PC stabilization might prevent irreversible microvasculature loss.

In addition to the CNS-mv-on-a-chip, we validated the obtained results, using microvascular drop model. The drop model reproduced key features of diabetic microvascular pathology, with both hyperglycemia and inflammation reducing vessel coverage area and the connectivity ratio (Fig. 6E, F). Inflammation (IT and GIT) caused an even stronger effect, reducing number of branches per mm^2^, and total vessel length. Notably, only the combined diabetic conditions (GIT) increased the average branch length, a phenotype consistently observed in both CNS-mv-on-a-chip (Fig. 4D) and microvascular drop models. The reproducibility across two platforms demonstrates the robustness of the diabetic microvascular pathology and highlights the microvascular drop as a cost-effective, easily scalable tool for high-throughput drug screenings. We have successfully allied the microvascular drop platform to assess *de novo*-designed VEGF inhibitors [58], further confirming its advantages for preclinical therapeutic evaluation.

The open geometry of the microvascular drop model, in contrast to enclosed design of microfluidic platforms, allows to directly quantify vessel sprouting. This proved particularly useful for assessing the effects of IL-6 on microvasculature morphology. While RNA-seq revealed upregulation of IL-6 under hyperglycemia (Fig. 3B), no morphological changes were observed in CNS-mv-on-a-chip (Fig. 4A, B). The microvascular drop platform revealed a strong pro-angiogenic effect of IL-6, marked by an increased vessel coverage, number of branches per mm², and branch length (Fig. 6E, F). Moreover, IL-6 promoted vessel sprouting (Fig. 6E, G). These findings align with previously shown data, demonstrating that elevated IL-6 in blood enhances vascular parameters, such as increased arterial diameter, enhanced capillary density, and improved blood flow [59]. In the brain and retinal setting, however, enhanced vessel sprouting might indicate the pathological neovascularization, and rather contribute to disease progression [60, 61], since newly sprouted blood vessels frequently can be associated with pathological morphology and high permeability.

To further enhance the physiological relevance of CNS-mv-on-a-chip model, we incorporated macroglial cell types – Müller glia cells (MGs), and astrocytes (ACs) – wich are key components of the neurovascular unit. In the brain, ACs provide structural support and regulate homeostasis, and barrier function [62]. Several self-organized CNS-mv models have successfully integrated ACs [22, 24, 63]. In contrast, much less is known about the role of MGs within retinal neurovascular unit. MGs are the primary glia type in the retina, with processes spanning the retinal thickness and contacting every cell type [64, 65]. Their processes ensheath capillaries, mirroring the tiled AC endfeet, wrapping brain capillaries. Although ACs and MGs share many morphological features, no side by side comparison of their roles in the CNS microvasculature has been reported to date as well as no microfluidic models, to our knowledge, incorporate MGs.

Both ACs and MGs were successfully integrated into the CNS-on-a-chip, extending their endfeet towards vessels. We observed several similarities between MG and AC effects on CNS-mv-on-a-chip morphology, including reduced vessel diameter, coverage area, and connectivity Fig. 7C, D). However, ACs appear to cause stronger effects, as reducing AC numbers in CNS-mv-on-a-chip mitigated these changes and increased similarities between MG and AC-induced effects. Additionally, MG integration promoted capillary network complexity, marked by increased total vessel length, and the number of branches per mm^2^ (Fig. 7D).

Functionally, MGs did not alter the apparent permeability coefficient, whereas AC integration significantly increased the vessel permeability, compromising barrier integrity (Fig. 7E, F). While ACs *in vivo* support the blood-brain-barrier integrity, hiPCS-derived ACs have been reported to disrupt vascular stability, likely due to their immature or reactive state. Previous studies have shown that flow through the vascular network can rescue this phenotype [24]. In our CNS-mv-on-a-chip the gravity-driven flow was applied, which may have been insufficient to promote AC maturation. In contrast, incorporation of hiPSC-derived MGs preserved barrier tightness, suggesting that EC-PC interactions in the baseline CNS-mv-on-a-chip model were sufficient for maintaining a restrictive barrier function.

We further established the iBRB-mv-on-a-chip as a first microfluidic model incorporating MGs and used it to model diabetic vascular pathologies. This is particularly significant since MG dysfunction under inflammatory conditions is one of the leading contributors to neovascularization and neural tissue damage [66]. The incorporation of MGs did not alter PC coverage of iBRB microvasculature (76.1 ± 6.5 % for EC:PC:MG vs. 75.5 ± 4.0 % for EC:PC microvasculature, respectively), indicating that MGs do not compete with PCs for EC contact *in vitro*. Instead, MGs integrated around both ECs and PCs, consistent with their *in vivo* organization [67]. Importantly, the iBRB-mv-on-a-chip recapitulated hallmark diabetic microvascular pathologies, including vascular regression, ghost vessel formation, and reduced PC coverage, consistently with the vascular changes observed in the CNS-mv-on-a-chip (Fig. 8A, B, D, E, F).

Interestingly, MGs behaved differently from PCs under diabetic conditions: while both hyperglycemia and inflammation reduced PC coverage, MG coverage was preserved or even enhanced under complete diabetic conditions (GIT) (Fig. 8G, H). This result suggests that diabetic conditions induced stress-associated behavior of MGs, leading to remodeling of the capillary architecture and an enhanced sheath. This finding is consistent with published data showing that diabetic conditions induce reactive gliosis in MG cells, leading to progressive and irreversible retinal damage [68]. These results further highlight the importance of targeting MGs in future therapeutic development. Overall, our findings demonstrate that the iBRB-mv-on-a-chip successfully models key features of diabetic vascular pathology in a tissue-specific context. MG integration enables the study of MG-vascular interactions, which are crucial for retinal diseases such as DR and reactive gliosis. This platform therefore provides a valuable tool for investigating mechanisms of microvascular dysfunction and for the development of targeted therapies.

In this study, we developed and validated three complementary *in vitro* models that together provide a comprehensive platform for understanding and treating diabetic microvascular pathologies. Each model addresses specific needs in the drug development pipeline: the microvascular drop system enables rapid, cost-effective screening of hundreds of compounds in 96-well formats; the CNS-mv-on-a-chip provides detailed mechanistic insights into vascular dynamics, including real-time visualization of ghost vessel formation and EC-PC interactions; and the iBRB-mv-on-a-chip offers the first tissue-specific platform incorporating Müller glia for retinal disease modeling.

## Materials and methods

### hiPSC culture

Three hiPSC lines were used in the project: TMOi001 [69] (female, reprogrammed from CD34-positive cord-blood-derived progenitor cells), INDBi001 [70] (male, reprogrammed from keratinocytes), and KOL2.1J [71] (male, reprogrammed from skin fibroblasts). Cells were cultured in a humidified incubator at 37 °C, 5 % CO_2_ and 5 % O_2_. For EC differentiation, hiPSCs were cultured on Geltrex-coated plates in E8 medium (Thermo Fisher Scientific, A1413301, A1517001); for PC differentiation hiPSCs were cultured on Matrigel-coated plates (Corning, CLS356234) in FTDA [72] medium. When hiPSCs reached 70-90 % confluency, they were passaged with Versene (Thermo Fisher, A15040066). mCherry- or GFP-labeled lines were obtained by transfecting the TMOi001 hiPSC line with lentiviral constructs, carrying fluorescent protein genes under the EF1α promoter (pLVX-EF1a-mCherry-N1 and pLVX-EF1α-AcGFP1-C1, Takara).

### Endothelial cell (EC) differentiation

hiPSCs were differentiated to ECs as described previously [26]. Briefly, hiPSCs were singularized with TrypLE (Thermo Fisher, A12604021) and 5 x 10^5^ cells were resuspended in 3 ml aggregation medium supplemented with 50 µM of Y-27632 (Stem Cell Technologies, 7232) to induce embryoid body (EB) formation. Mesodermal specification was induced by transferring EBs to N2B27 medium supplemented with 12 µM CHIR99021 (Selleckchem, S2924) and 30-60 ng/ml BMP-4 (depending on the cell line) (Peprotech, 120-05). Three days later the vascular specification was induced by supplementing the medium with 100 ng/ml VEGF-A (Peprotech, 100-20) and 2 µM Forskolin (Sigma-Aldrich, F6886). On day 5, cells were plated on 0.2 % gelatin-coated plates and cultured in StemPro-34 medium (Thermo Fisher, 10639011) supplemented with 100 ng/ml VEGF-A and 100 ng/ml FGF-2 (Peprotech, HZ-1038, 100-18B). On day 10, cells were sorted by CD31 expression using a MicroBead Kit (Miltenyi Biotec, 130-091-935), according to the manufacturer’s instructions. CD31^+^ ECs were cultured on 0.2 % gelatin-coated plates in MV medium (PromoCell, C-22020) that contains physiological (5.5 mM) glucose level supplemented with 30 ng/ml VEGF-A and 20 ng/ml FGF-2. Freshly sorted ECs were considered passage 0 (P0); for microvasculature formation ECs P2-3 were used.

### Pericyte (PC) differentiation

PCs were differentiated from neural crest stem cells (NCSCs) using a previously published protocol [27]. Briefly, the NCSC differentiation was induced by culturing singularized hiPSCs on GFR-Matrigel-coated (Corning, CLS356231) plates in E6-CSFD medium. On day 15 of differentiation, NCSCs were sorted using MicroBead Kit (Miltenyi Biotec, 130-097-127). PC differentiation was induced by culturing cells on uncoated plates in E6 medium supplied with 10 % FBS for at least 7 days. PCs were frozen or used immediately.

### Glia cell differentiation

Astrocytes (ACs) were differentiated as previously described [73]. EBs were formed in a 96-well plate in mTeSR medium (StemCell Technologies, 100-1130) supplemented with 10 µM blebbistatin (Sigma-Aldrich, TA9H97F31488) and 10 µM Y-27632 (Stem Cell Technologies, Y-27632). The following day, the medium was replaced with N2 medium (DMEM/F-12 with Hormonmix: 88 µM Putrescin, 20 µg/ml insulin, 80 µg/ml holotransferrin, 16 nM progesterone, 24 nM sodium selenite). On day 7, EBs were transferred to a GFR-Matrigel-coated plate in N2 medium. On day 15, neural rosettes were detached. On day 21, the medium was changed to N2 medium containing 10 ng/ml EGF (Peprotech, HZ-1326) and 10 ng/ml FGF-2, and cultures were maintained until day 120. At day 120, ACs were dissociated into single cells using Accutase (Sigma-Aldrich, A6964) and plated on fibronectin-coated plates (5 µg/cm²) (Corning, CLS354008) in N2 medium supplemented with 10 ng/ml EGF and 10 ng/ml FGF-2. Maturation was induced by switching to N2 medium containing 10 ng/ml BMP-4 (Peprotech, HZ-1128) and 10 ng/ml CNTF (Peprotech, HZ-1331) for 7 days.

MGs were isolated from retinal organoids (ROs) as previously [74]. ROs were differentiated based on a protocol published by Zhong et al. [75]. ROs at 125-225 days of differentiation were dissociated with the Neurosphere Dissociation Kit (Miltenyi Biotec, 130-095-943) and seeded onto fibronectin-coated plates (5 µg/cm²) in DMEM medium containing 10 % FBS (Thermo Fisher, A5256801), 1 % 100x non-essential amino acids (Thermo Fisher,11140) and 1 % 100x GlutaMAX (Thermo Fisher, 35050), supplemented with 40 ng/ml EGF (Peprotech, AF-100-15-500UG) and 40 ng/ml FGF-2 (Peprotech, AF-100-18B-50UG). Cells isolated from ROs were considered P0 and propagated up to P4. Prior to loading onto microfluidic chips, MGs were matured by growing them to 90 % confluence and then culturing them in medium without EGF and FGF-2 for 7 days.

### Microvasculature formation on the chip

For CNS-mv formation, ECs and PCs were detached and mixed in a 1:1 ratio with a total cell concentration of 1.6 x 10^7^ cells/ml (80,000 cells per chip channel) in a 5 mg/ml fibrinogen solution (Sigma-Aldrich, F3879) with 4 U/ml thrombin (Sigma Aldrich, T4648). 10 µl of this cell suspension was seeded in the middle channel of a microfluidic chip (Aimbiotech, DAX-1). After 30 min gel polymerization at room temperature, MV medium with 30 ng/ml VEGF-A and 20 ng/ml FGF-2 was added to side channels, inducing a gravity flow (90 µl of medium was added to the top channel ports and 50 µl to the bottom ports). For triple cell culture, glial cells were added in 3:3:1 or 5:5:1 EC:PC:glia ratio, with final cell concentration 80,000 cells per chip. The chips were cultured for seven days with daily media changes. To induce vasculature maturation, on day four, VEGF-A was removed from the medium. To model hyperglycemia, glucose (Carl Roth, 10358434) was added to MV medium to the final concentration of 25 mM, 50 mM, or 75 mM. As corresponding osmotic controls, we used MV medium with mannitol (19.5 mM, 44.5 mM, or 69.5 mM) (Sigma Aldrich, M4125). An inflammatory profile was induced by adding (1) 1 ng/ml TNF-α (Peprotech, HZ-1014), (2) 1 ng/ml IL-1β (Peprotech, HZ-1164), (3) 1 ng/ml IL-6 (Peprotech, 200-06), (4) TNF-α and IL-1β combination (IT). To induce complete diabetic conditions (GIT) we used MV medium with 25 mM glucose supplemented with 1 ng/ml TNF-α and 1 ng/ml IL-1β. For inflammatory cytokines treatments mock controls were used.

### Microvasculature formation on the drop

The procedure for drops loading was similar to the chip loading described above, with slight modifications: the total cell number per one drop was 6 x 10^6^ cells/ml suspended in 5 µl of fibrin gel. Drops were cultured on non-tissue treated 48- or 96-well plates or on glass cover slips. Medium was changed daily.

### Immunostaining

Cells, 3D drops, or microfluidic chips were fixed with 4 % PFA for 20 minutes, incubated in blocking solution (10 % normal donkey serum containing 0.2 % Triton-X-100) for 1 h at room temperature, then incubated with primary antibodies (CD31: Abcam, ab9498; PDGFRβ: Cell Signalling Technology, 3169; Claudin-5: Invitrogen, 35-2500; VE-cadherin: Cell Signalling Technology, 2158; ZO-1: Thermo Fisher Scientific, 33-9100; Phalloidin: Invitrogen, A22287; CD146: Biolegend, 361001; CD34: Abcam, ab81289; NG2: Chemicon, ab5320; SMA: Abcam, ab5694; Collagen IV: Chemicon, AB769; Vimentin: Abcam, ab73159; Nestin: Abcam, ab24692-50, ERG −1/2/3: Santa Cruz Biotechnologies, sc-376293) in blocking solution overnight at 4 °C, and then with secondary antibodies (Donkey anti-Mouse IgG Secondary Antibody Alexa Fluor 488 polyclonal, R37114; Donkey anti-Rabbit IgG Secondary Antibody Alexa Fluor 568 polyclonal, A10042; Donkey anti-Goat IgG Secondary Antibody Alexa Fluor 647 polyclonalm A32849 from Thermo Fisher) diluted in blocking solution for 1 h at room temperature. Nuclei were stained with 10 µg/ml Hoechst-33342 (Sigma-Aldrich, 14533) for 20 min.

### Cryosectioning

Microvasculature on drops were fixed in 4 % PFA solution for 30 min at room temperature, detached from the growing plate and embedded in Tissue Tek® O.C.T^TM^ (Sakura, 4583) in a cryomold. The frozen block was cut at −20 °C with 14 µm slice thickness.

### Microvasculature morphology analysis

Fluorescent images of chips and drops were taken and analyzed using ImageJ. The signals of mCherry- or, GFP-labeled cells, or cells stained with CD31 antibodies, were used to identify ECs. Images were binarized, outliers were removed, and holes were closed. Skeletonization was performed using the Skeletonize2D/3D [76] plugin, and skeleton parameters were measured. Skeleton parameters were used to calculate total vessel length, average branch length, number of branches and the connectivity ratio (number of junctions divided by the number of endpoints). The total area and vessel coverage area fraction were measured from the binarized image. The average lateral vessel diameter was estimated by dividing the total vascular coverage area by the total vessel length. To calculate vascular sprouting from drops, the central part of the drop was removed, and only the outside rim was retained for the skeletonization analysis to quantify the total vessel length and the number of branches.

### Pericyte coverage and ghost vessel quantification

For PC coverage analysis, z-stack images with slices of 1.5 µm for the entire thickness of the chip were acquired. The average intensity z-projection was used to identify the vasculature area positive for ECs. A median filter, followed by thresholding, was applied to binarize images. Similarly, the PC area was defined. The overlap of vascular area and PC area was defined as PC-covered area. The PC-covered area divided by the vascular area was used as the PC coverage fraction. For ghost vessel analysis, the binarized image of vascular area was subtracted from the collagen IV positive stained area to obtain the remaining ghost vessel area. The ghost vessel fraction was quantified as the ghost vessel area divided by vascular area.

### Permeability assay

For the permeability assay, the bottom side channel was seeded with ECs to ensure the connection of the microvasculature to the media channel. To this end, the side channel was coated with 0.2 % gelatin for 1 h immediately after middle channel polymerization. Then, the coating solution was washed with fresh MV medium, and on day 2, 5 µl (2 x 10^6^ cells/ml) of ECs suspension was loaded into the side channel and incubated for 1.5 h to ensure EC attachment. Then, the channel was washed with MV medium and cultured as described above. To demonstrate the perfusion of the microvascular lumen by cells, 7.5 µm compensation beads were used. The Alexa-488-labeled beads were loaded in the top channel, and the chip was immediately imaged. To quantify microvascular permeability, the side channel was loaded with 0.1 mg/ml 70 kDa FITC-dextran (Sigma-Aldrich, FD10S-100MG) and 0.05 mg/ml 10 kDa 640R-Dextran (Biotium, 80115). Images were taken immediately after loading the side channel and then every 30 s for 10 min. Image analysis was performed with ImageJ: the microvasculature was detected based on the mCherry signal of labeled ECs; the region was binarized, outliers were removed, and holes were closed; mean fluorescence intensity (MFI) of dextran signals was measured inside and outside the vasculature; the vascular region was skeletonized, and the vessel perimeter was calculated. The apparent permeability coefficient was calculated as described previously [22, 77].

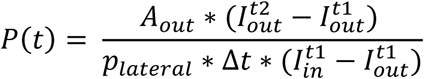

Here, 𝐴_𝑜𝑢𝑡_ is the area of the intervascular space, 𝐼 the MFIs of the dextran signal inside and outside the vascular area at time points 1 and 2, Δ𝑡 = 𝑡2 − 𝑡1 is the time interval, 𝑝_𝑙𝑎𝑡𝑒𝑟𝑎𝑙_ is the calculated vessel perimeter.

### RNA sequencing analysis

Cells were collected from the chips (four independent replicates for each condition, with two chips pooled to increase RNA yield), and RNA was isolated using the RNeasy Micro Kit (Qiagen, 74004) according to the manufacturer’s protocol. RNA sequencing (RNA-seq) was performed by the Quantitative Biology Center (QBiC), Institute for Medical Genetics and Applied Genomics (IMGAG) and Institute for Medical Microbiology and Hygiene (MGM) of the University of Tübingen. Quality and quantity of RNA were assessed by fluorescence-based quantification and RNA integrity analysis to ensure high quality RNA. Sequencing was performed with the Illumina NovaSeq 6000. QBiC also provided primary data analysis. FASTQ files were assessed for quality with FastQC [78], trimming was performed with TrimGalore [79], ribosomal RNA was removed using SortMeRNA [80], and reads were aligned to the human genome using STAR [81]. FeatureCounts [82] and Salmon [83] were used to obtain gene and transcript counts. The RNA-seq dataset is available under accession number GSE 300952. RNA-seq data were analyzed using the DESeq2 package for R [84]. The bioinformatical analysis included the differential gene expression analysis (DGE) and gene enrichment analysis (GEA). For GEA, a cutoff of log2FC > |1| and adjusted P-value < 0.01 was used. Pathway enrichment analysis with the Hallmark set the KEGG was done with the clusterProfiler package [85]. DGE results were visualized by volcano plot (padj < 0.05, Log2FC >1) via ggplot2 package [86] and heatmap (padj<0.1, Log2FC>1, baseMean >= 20) by using pheatmap package [87].

### qPCR analysis

RNA was isolated from MACS-sorted ECs and PCs from the CNS-mv-on-a-chip using the Quick Extract RNA Extraction Kit (Biozym, 101061). Genomic DNA was removed with RNase-free DNase I, and cDNA was synthesized using the GoTaq 2-Step RT-qPCR System. RT-qPCR analysis was performed using a high-throughput Fluidigm system (BioMark). First, a pre-amplification step with TaqMan primers (Thermo Fisher Scientific) was carried out for GAPDH (Hs99999905_m1), HMBS (Hs00609297_m1), IL1B (Hs01555410_m1), JUN (Hs01103582_s1), and NFKBIA (Hs00355671_g1). Pre-amplified samples were then used for the Fluidigm run. For each gene and sample, CT values from four technical replicates were averaged. GAPDH and HMBS were used as housekeeping genes, and their mean CT value per sample was subtracted from the CT value of each target gene to obtain the ΔCT value. Relative expression was calculated using the 2^-ΔΔCT^ method.

### Microscopy

For fluorescence and phase-contrast images Axio Observer microscope (Carl Zeiss, Germany) was used with Apotome 3, EC Plan-Neofluar 10x and 20x Objectives, 545-565, 583-601, 545-565, 623-670 filters, and X-Cite Xylis Lamp.

### Statistical analysis

Results are presented as mean with standard deviation (SD). Visualization and statistical analysis were done with GraphPad Prism. One-way Analysis of Variance (ANOVA) was done to evaluate statistical significance. For some analysis treatments were normalized to the control of each independent experiment and one sample t-test was performed to evaluate statistical significance. For each graphs N and n are given, where N depicts the number of independent experiments and n number of samples in total. For all analysis, statistical significance was represented as: **** p<0.0001, *** p<0.001, ** p<0.01, * p>0.05.

## Acknowledgments

Institutional Review Board Statement

All procedures were in accordance with the Helsinki Convention and approved by the Ethical Committee of the Eberhard Karls University Tübingen (no. 396/2021BO2).

## Funding

This research was funded by the Fortüne program of the University of Tübingen, Germany (2667-0-0 to N.P.).

## Supplementary materials

**Supplementary Figure 1.**
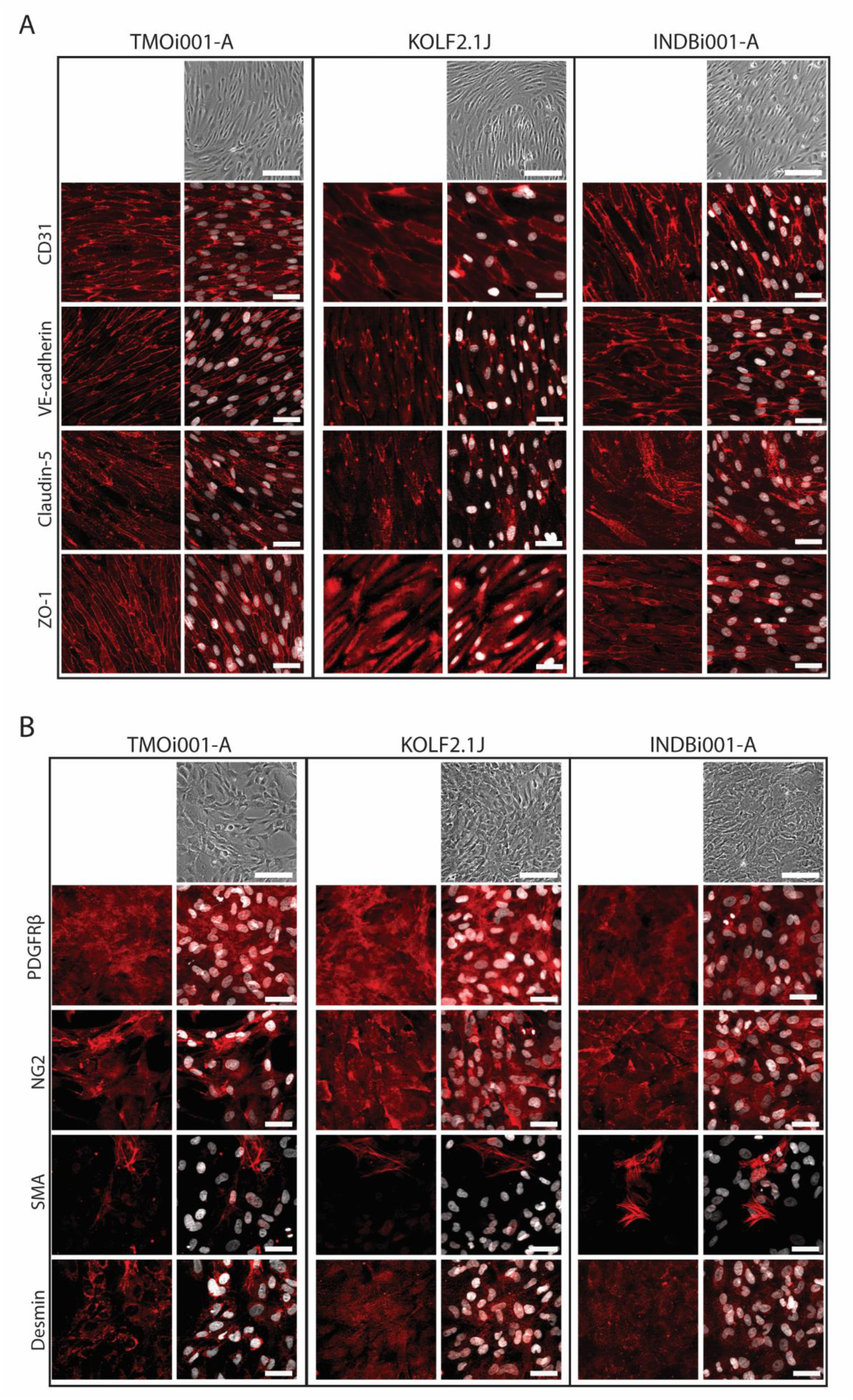
**Characterization of hiPSC-derived ECs and PCs via immunocytochemistry.** (A) Brightfield images of EC from three different hiPSC cell lines, TMOi001-A, KOLF2.1J and INDBi001-A. Representative immunofluorescence images of EC marker CD31, tight junction and adherens junction marker VE-cadherin, claudin-5 and ZO-1. Scale bar: Brightfield images 200 µm, fluorescence images 50 µm. (B) Brightfield images of PC from three different hiPSC cell lines, TMOi001-A, KOLF2.1J and INDBi001-A. Representative immunofluorescence images of PC marker PDGFRβ, NG2, SMA and Desmin. Scale bar: Brightfield images 200 µm, fluorescence images 50 µm.

**Supplementary Figure 2.**
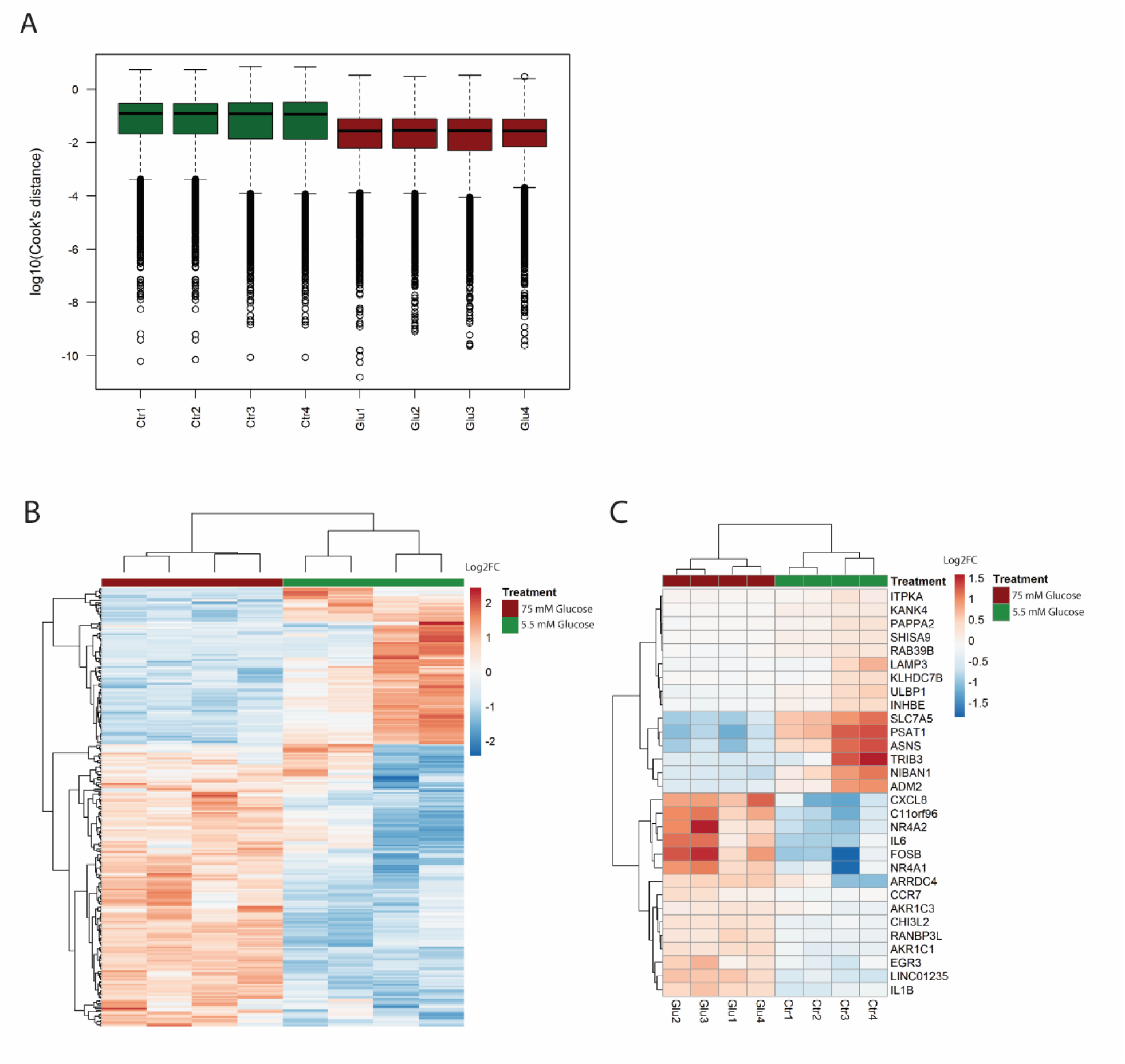
**RNAseq analysis of diabetic conditions.** (A) Quality control analysis included cook’s distance analysis to identify possible outlier and confirm a robust model fitting. (B) Heatmap shows all differentially expressed genes clustered by gene expression profile for each sample. 5.5 mM glucose and 75 mM glucose show similar clustering.(C) Heatmap with the 30 most significant up and down regulated genes.

**Supplementary Figure 3.**
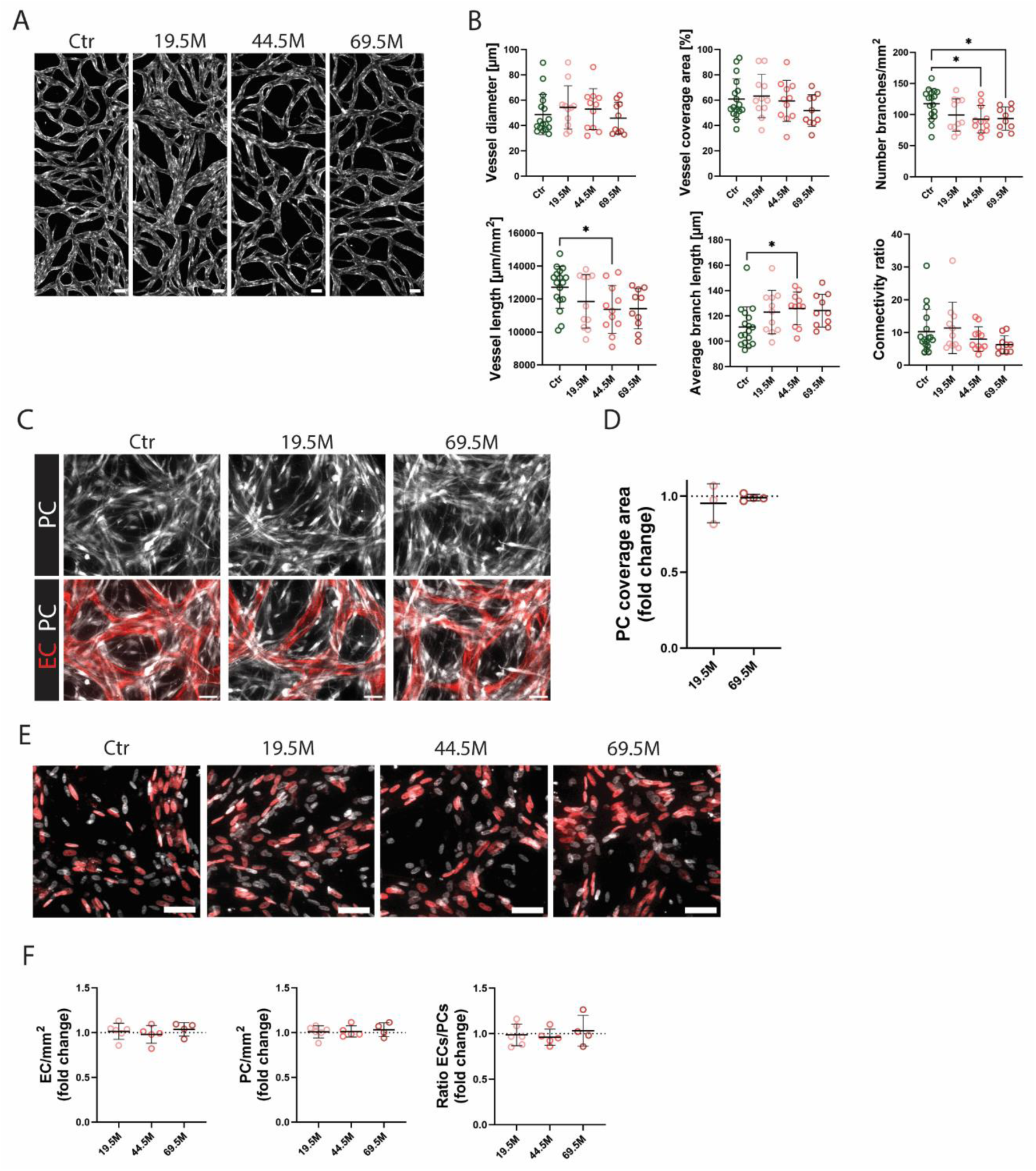
**Osmotic control to hyperglycemic treatments in the CNV-mv-on-a-chip model.** (A) Fluorescence images of mCherry-labeled ECs in the CNS-mv-on-a-chip. Osmotic control included 19.5 mM, 44.5 mM and 69.5 mM mannitol. Scale bar 100 µm. (B) Quantification of microvascular parameters of osmotic controls. (C) Fluorescence images of mCherry-labeled ECs (red) and GFP-labeled PCs (white) cultured in control, 19.5 mM or 69.5 mM mannitol. Scale bar 50 µm. (D) Quantification of PC coverage exposed to osmotic pressure controls. Values are normalized to control group in each independent experiment. (E) Fluorescence images of CNS-mv-on-a-chips stained with ERG-1/2/3 (red) to identify EC nuclei and Hoechst (white). Scale bar 50 µm. (F) Quantification of EC and PC number and EC/PC ratio in osmotic controls. Values are normalized to control group in each independent experiment.

**Supplementary Figure 4.**
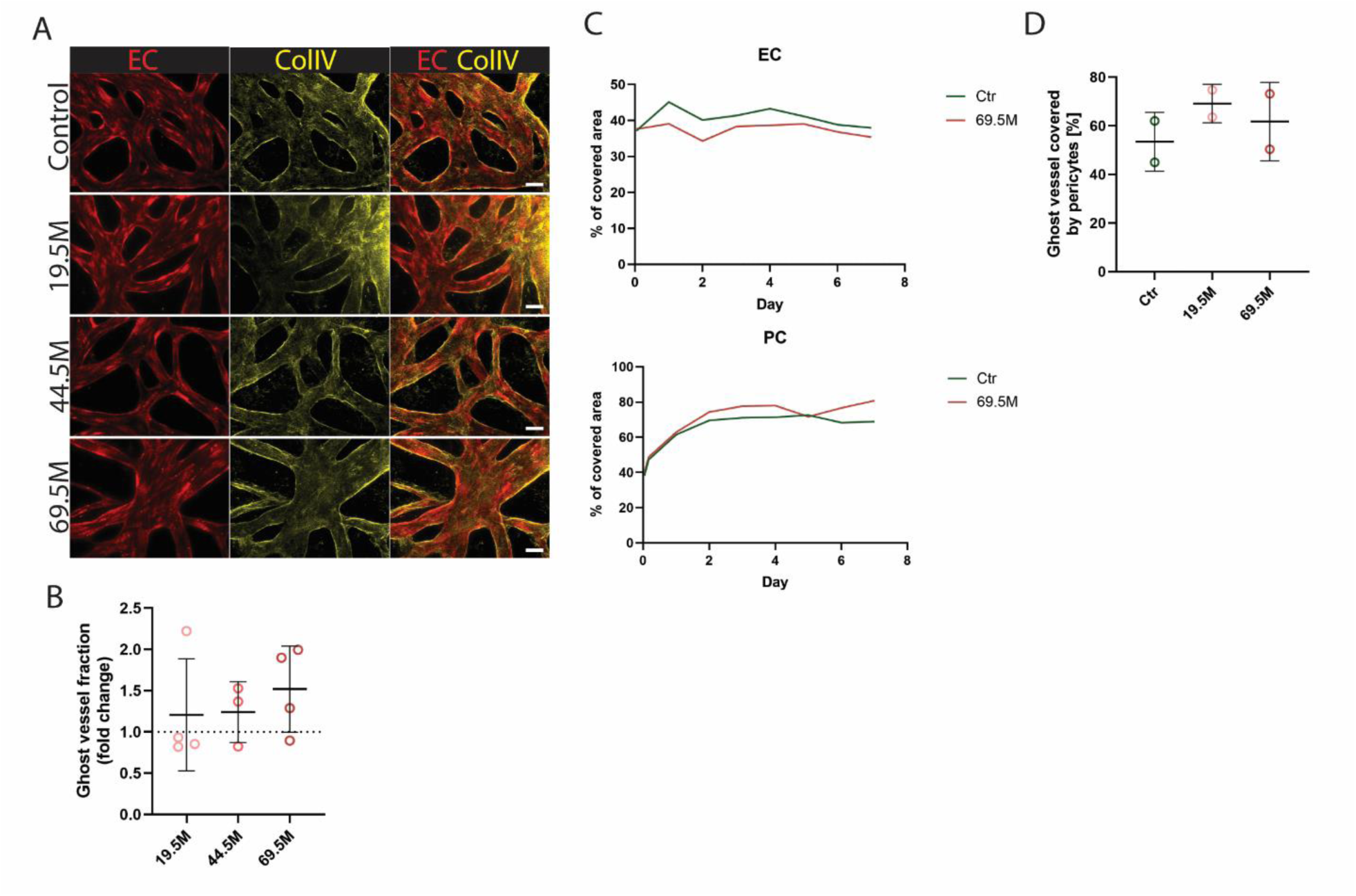
**The osmotic controls for ghost vessel formation.** (A) Fluorescence images CNS-mv-on-a-chips with stained collagen IV and mCherry-labeled ECs. Chips were exposed to osmotic pressure controls with 19.5 mM, 44.5 mM and 69.5 mM mannitol. Scale bar 50 µm. (B) Quantification of ghost vessel fraction in microvasculatures exposed to osmotic pressure controls. (C) Quantification of area fraction covered by ECs and PC over 7 days under osmotic controls. (D) Quantification of ghost vessels covered by PCs in CNS-mv-on-a-chips exposed to osmotic controls.

**Supplementary Figure 5.**
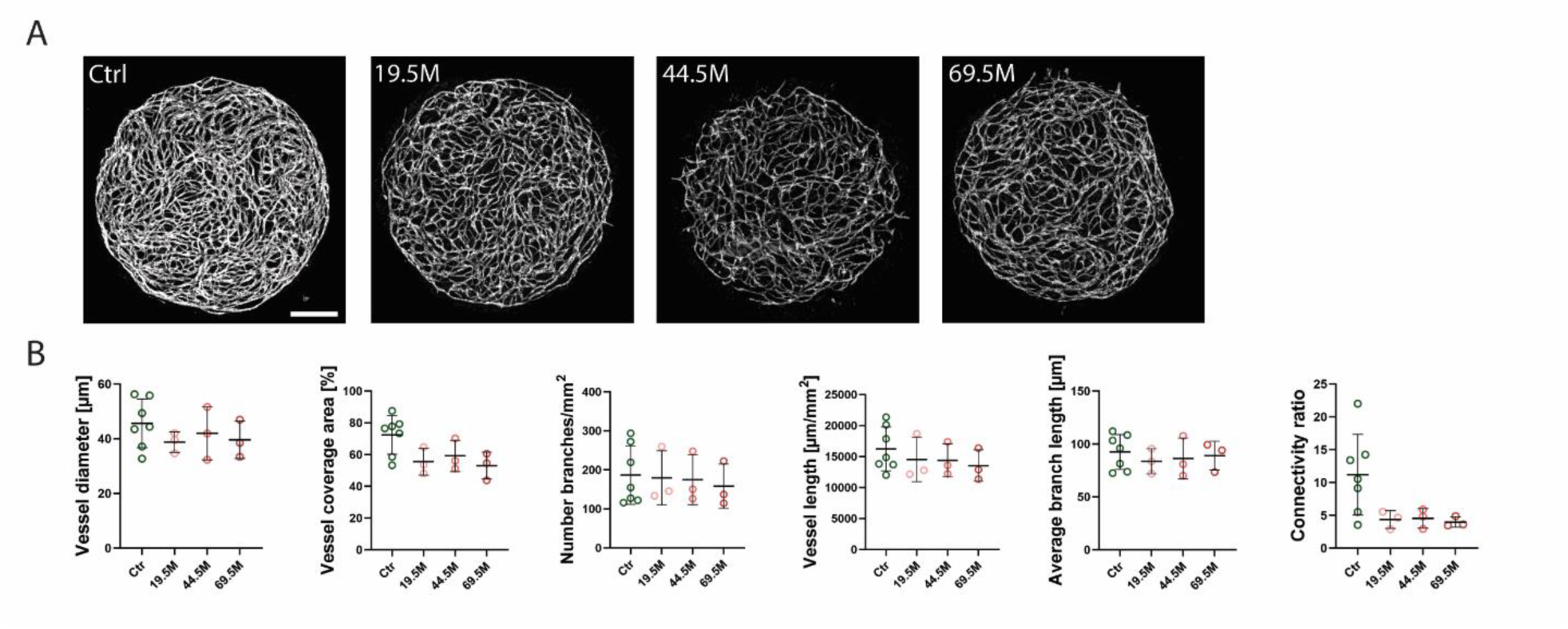
**Osmotic control to hyperglycemic treatments in the microvascular drop model.** (A) Fluorescence images of mCherry-labeled ECs. Osmotic control included 19.5 mM, 44.5 mM and 69.5 mM mannitol. Scale bar 500 µm. (B) Quantification of microvascular parameters of osmotic controls.

**Supplementary Figure 6:**
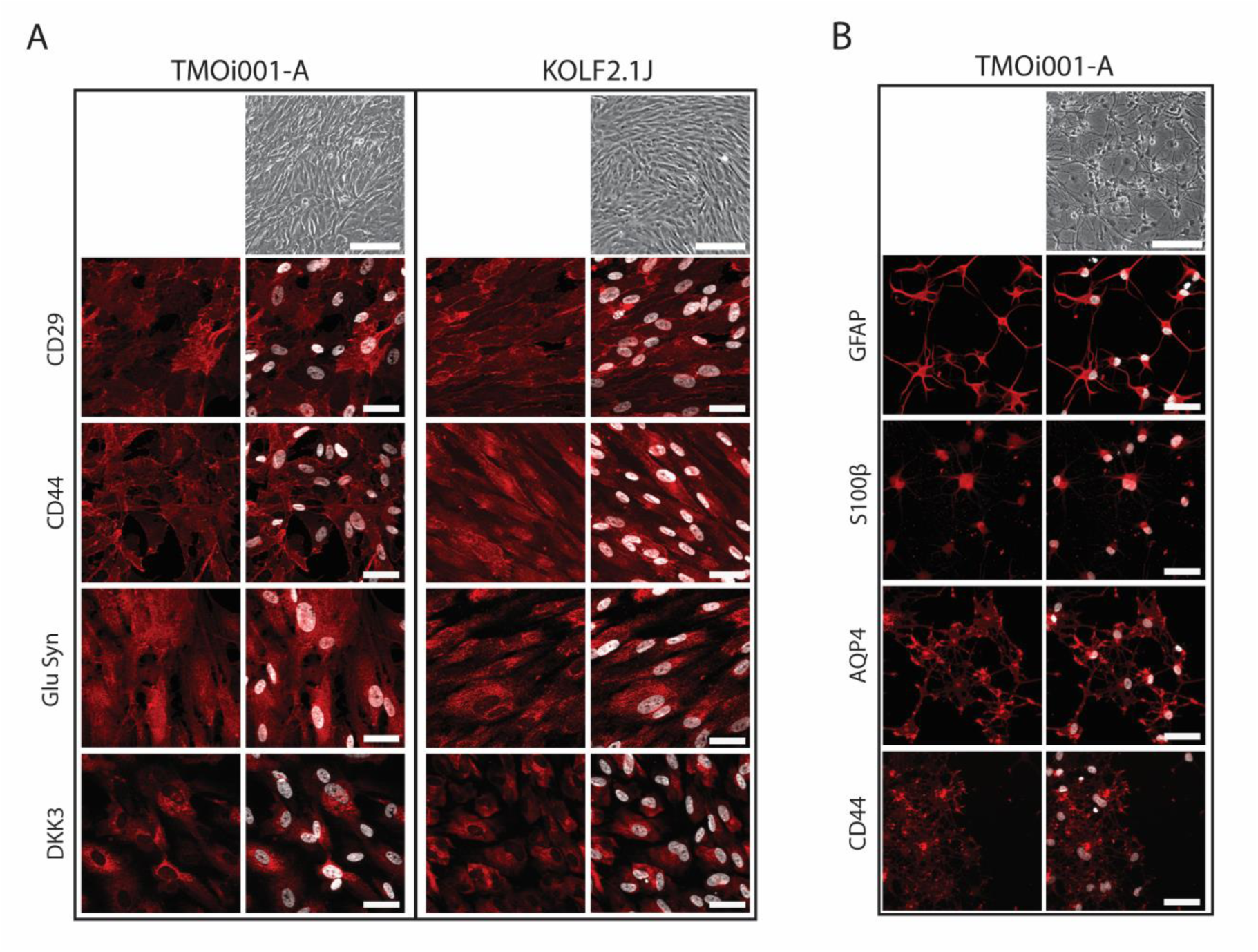
**Characterization of hiPSC-derived MGs and ACs via immunocytochemistry.** (A) Brightfield images of MGs from two different hiPSC cell lines, TMOi001-A and KOLF2.1J. Representative immunofluorescence images of MG marker CD29, CD44, GluSyn and DKK3. Scale bar: Brightfield images 200 µm, fluorescence images 50 µm. (B) Brightfield images of AC from one hiPSC cell line, TMOi001-A. Representative immunofluorescence images of AC marker GFAP, S100β, AQP4 and CD44. Scale bar: Brightfield images 200 µm, fluorescence images 50 µm.

**Supplementary Figure 7.**
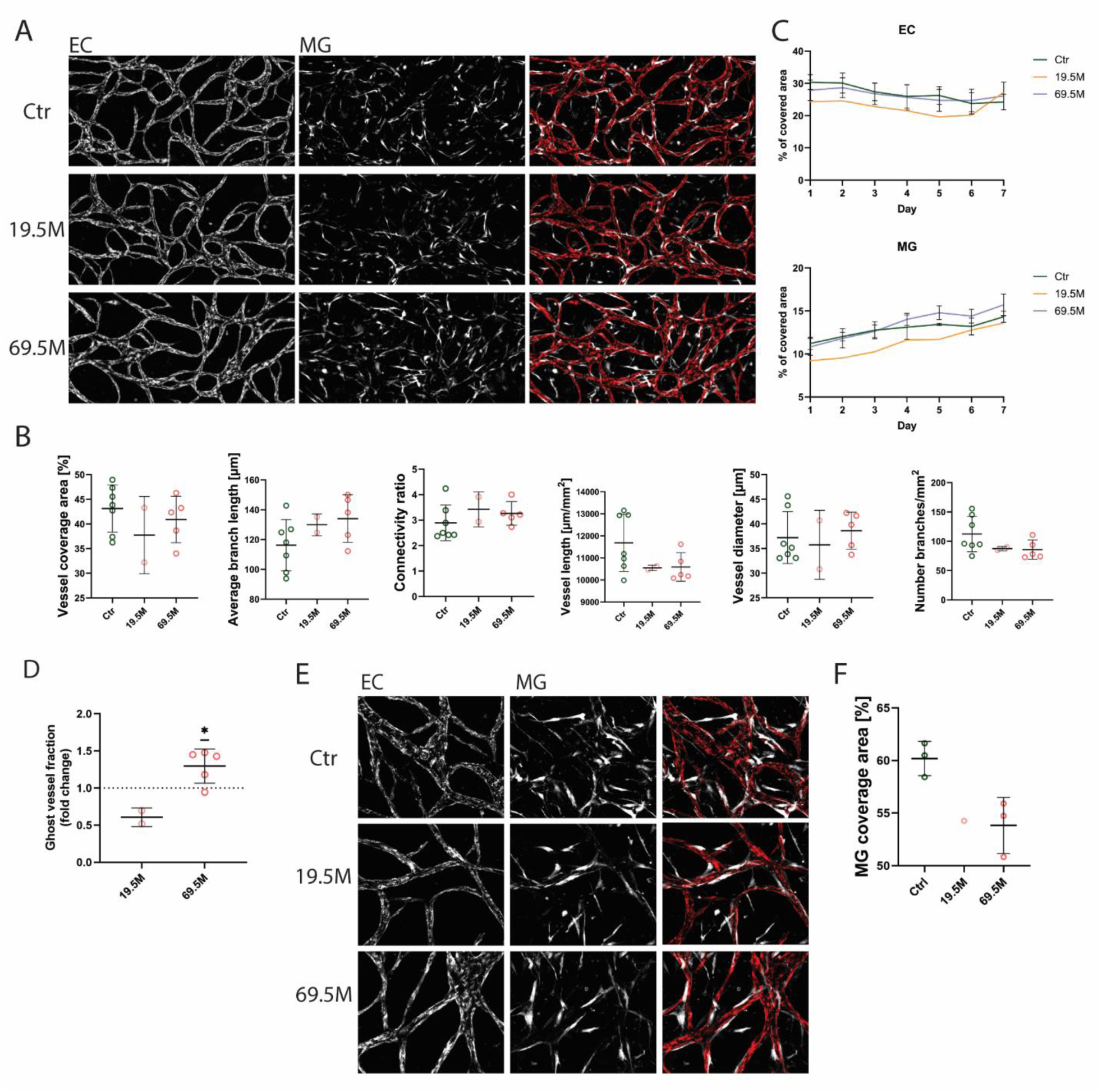
**Osmotic control to hyperglycemic treatments in the iBRB-on-a-chip model.** (B) Fluorescence images of mCherry-labeled ECs in the iBRB-on-a-chip. Osmotic control included 19.5 mM, 44.5 mM and 69.5 mM mannitol. Scale bar 100 µm. (C) Quantification of microvascular parameters of osmotic controls. (D) Quantification of the area fraction covered by ECs and MGs over 7 days under osmotic controls. (E) Quantification of the ghost vessel fraction under osmotic controls.

## Notes

### Competing Interest Statement

The authors have declared no competing interest.

### Summary of Updates

The author list order was corrected

